# Does the rainbow trout ovarian fluid guide the spermatozoon on its way to the egg?

**DOI:** 10.1101/2021.08.05.453951

**Authors:** Vitaliy Kholodnyy, Borys Dzyuba, Marek Rodina, Hermes Bloomfield-Gadêlha, Manabu Yoshida, Jacky Cosson, Sergii Boryshpolets

## Abstract

Fertilization of freshwater fish occurs in an environment that may negatively affect the gametes; therefore, the specific mechanisms triggering the encounters of gametes would be highly expedient. The egg and ovarian fluid (OF) are likely the primary sources of these triggers in fish that we confirmed here for rainbow trout *Oncorhynchus mykiss*. The ovarian fluid significantly affected the spermatozoa performance: it supported high velocity for a more extended period and changed the motility pattern from tumbling in water to straightforward moving in the ovarian fluid. Rainbow trout OF induced a trapping chemotaxis-like effect on activated male gametes, and this effect depended on the properties of the activating media. The interaction of the spermatozoa with the attracting agents was accompanied by their “turn-and-run” behavior involving asymmetric flagellar beating and Ca^2+^ concentration bursts in the bent flagella segment, characteristic of the chemotactic response. Collectively, the ovarian fluid creates the optimal environment for rainbow trout spermatozoa performance, being an effective promoter of fertilization. The individual peculiarities of the egg (ovarian fluid) – sperm interaction in rainbow trout reflect the specific features of the spawning process in this species.

## Introduction

Sexual reproduction grounds on the fertilization of female ova by male spermatozoa. For successful fertilization, the spermatozoa must reach the egg in due time after ovulation. Externally fertilizing freshwater fish spermatozoa are activated after direct contact with freshwater, and its motility is only sustained for a very limited period, typically tens of seconds [1]. Freshwater fish eggs are relatively large comparing, *e.g.,* to mammal oocytes and possess a specialized fertilization site, namely, a micropyle, which is to be found by the diminutive sperm cell during its short motility period in an environment that is constantly moving and changing. Under these conditions, reproductive success is chronically limited to the ability of spermatozoa to find the egg and reach the fertilization site, and thus they depend on mechanisms that increase the probability of sperm-egg encounter.

What are these mechanisms based on? Generally, ideas on this matter vary from the predominance of a “male factor”, *e.g.*, “fair raffle” concept, w hen the success of a particular male depends only on the number of spermatozoa it could provide to the “fertilization lottery” [2], to the existence of a specific female post-mating control, *e.g.*, “cryptic female choice” [3]. And finally, the combination of many factors, including environmental ones, is considered, *e.g*., guidance hypothesis [4]. There are numerous assumptions that the gametes’ encounter is controlled by stimuli or signaling which come from the egg or accompanying female fluids, and confirmations were found across many taxa, including marine invertebrates, mammals, and in a few fish species [5, 6]. These stimuli may affect the behavior of sperm cells, rendering both chemokinetic and chemotactic effects, and finally, the outcome of fertilization [7].

First observations on sperm activation and the rise in its motility traits in the vicinity of the eggs were made in marine invertebrates about 100 years ago [8], and “egg jelly”, a substance surrounding the eggs, was believed to be responsible for that. Marine invertebrates remain the main participants of the studies on the interaction of gametes, including the chemotactic issues, in particular, the most popular and complete spermatozoon chemotaxis models are based on sea urchins [9] and ascidians[10]. In fish, such a female effect provider can be the ovarian fluid (OF), which bathes mature fish oocytes in the ovarian cavity [11] and surrounds the eggs of many externally fertilizing female fish during the release of the spawned eggs through the oviducts into the water [12]. The composition of the OF varies among fish species, generally, it contains ions and different substances of protein nature, sugars, and lipids in different ratios [13–15].

There are several reports about the existence of chemokinetic reactions of fish male gametes, *i.e.,* changes in swimming velocity and trajectories of spermatozoa depending on the presence of ovarian fluid in the media [16–19] scrutinized recently by Myers et al. [20] in their meta-analysis. Interestingly, most of the studies presented in this analysis were done in salmonids, which is not surprising, considering their popularity as a research object for fish reproduction studies, and preconditioned by its wide usage in aquaculture and consequently high market value. The meta-analysis of Myers et al. [20] showed the overall enhancing effect of ovarian fluid on the velocity of the salmonid spermatozoa, pointing, however, to the high heterogeneity of the data. Nevertheless, there is still very little information beyond these kinetic observations. It is not clear if the ovarian fluid of salmonid fishes provokes any chemotactic effect or how it may trigger the success of egg fertilization, and there is no clear understanding of the mechanisms of motility enhancement provided by ovarian fluid.

In our study, we aimed to perform comprehensive testing of the interaction of spermatozoa and ovarian fluid, as well as to find the evidence or absence of chemotactic behavior in one of the representative of freshwater spawning fishes of the Salmonidae family, the rainbow trout *Oncorhynchus mykiss*, and to clear up the potential mechanism of gametes’ encounter guidance in this species.

To reach our goal, we resolved the following issues:

- How the presence of ovarian fluid affects the outcome of *in vitro* fertilization of eggs?
- What is the effect of ovarian fluid in the activation medium on the spermatozoa motility traits, including velocity and linearity of motion?
- Whether the ovarian fluid has a chemotactic effect on spermatozoa?
- Are the changes in spermatozoa motility associated with the specific agents in the ovarian fluid, including osmolarity and Ca^2+^ ions?

## Results

### Ovarian fluid enhances the in vitro fertilization outcome

It was already shown that the presence of ovarian fluid increases the success of *in vitro* fertilization in salmonids, *e.g.,* Caspian brown trout, *Salmo trutta caspius* [21]. We have performed our *in vitro* fertilization test to confirm this effect in the chosen fish model, *i.e., O. mykiss*. Moreover, we have considered the potential variability of the effect depending on the relatedness of the mates, in particular, if they belong to different populations. It was reported, *e.g.*, in guppy *Poecilia reticulata* [22] or lake trout *Salvelinus namaycush* [23]. In our study, one population was represented by albino males and females (originating from one long-term broodstock), and the other population consisted of regularly colored males (originating from one broodstock). The albino type is non-dominant, *i.e.*, fertilization of the egg from an albino female with the sperm from a regularly colored male will result in a regular color embryo/larva that allows a simple assessment of the sire.

It appeared during the preliminary motility tests that sperm from albino males had low motility in water, only up to 20%; however, the motility in the ovarian fluid was higher than 95%. Sperm from regularly colored males had more than 50% motility in water and more than 95% in the ovarian fluid. We have used two concentrations of spermatozoa in the activation medium, adding 0.5 or 5 µl of sperm to 8 ml of activating medium. Fertilization of the intact (non-washed) eggs in water with a lower concentration of spermatozoa (0.5 µl of sperm, *i.e.*, 150,000 spermatozoa per egg) resulted in a relatively low developing embryo count, less than 20% in the case of albino males, and around 40% in normal males (Figure 1a). In the case of mixed sperm, the outcome was about 20% fertilized eggs, and only 16% of them were albino embryos. Washing of the eggs to remove ovarian fluid and following fertilization in water resulted in a significant rise of fertilization with sperm from normal males (p<0.0005, binomial test with p-value corrected for multiple comparisons) and no significant changes for the albino male group (p=0.01). The amount of fertilized eggs in the mixed group rose as well (p<0.0005), the share of albino embryos stayed the same (p=0.28), nevertheless, their absolute number was higher compared to the case with intact eggs. The fertilization in the isotonic NaCl solution resulted in higher amount of developing embryos for albino males (p<0.0005), non-significant rise for normal color males (p=0.002) and a decrease in the mixed group (p<0.0005) compared to the fertilization in the water of washed eggs. The use of 100% ovarian fluid as an activation medium resulted in almost 100% success of egg fertilization in all cases, either in albino, normal males, or the mixed group. The share of albino embryos in the mixed group almost reached the half of total embryo count (p=0.002). Using of higher concentration of spermatozoa (adding of 5 µl sperm into the activation medium, i.e. 1,500,000 spermatozoa per egg) resulted in a high fertilization outcome in all cases with no great differences between the treatments (Figure 1b). Nevertheless the share of albino embryos in the group with fertilization in OF was again the highest comparing to other treatments and did not significantly differed from expected “ideal” 50% division (p=0.12).

**Figure 1.**
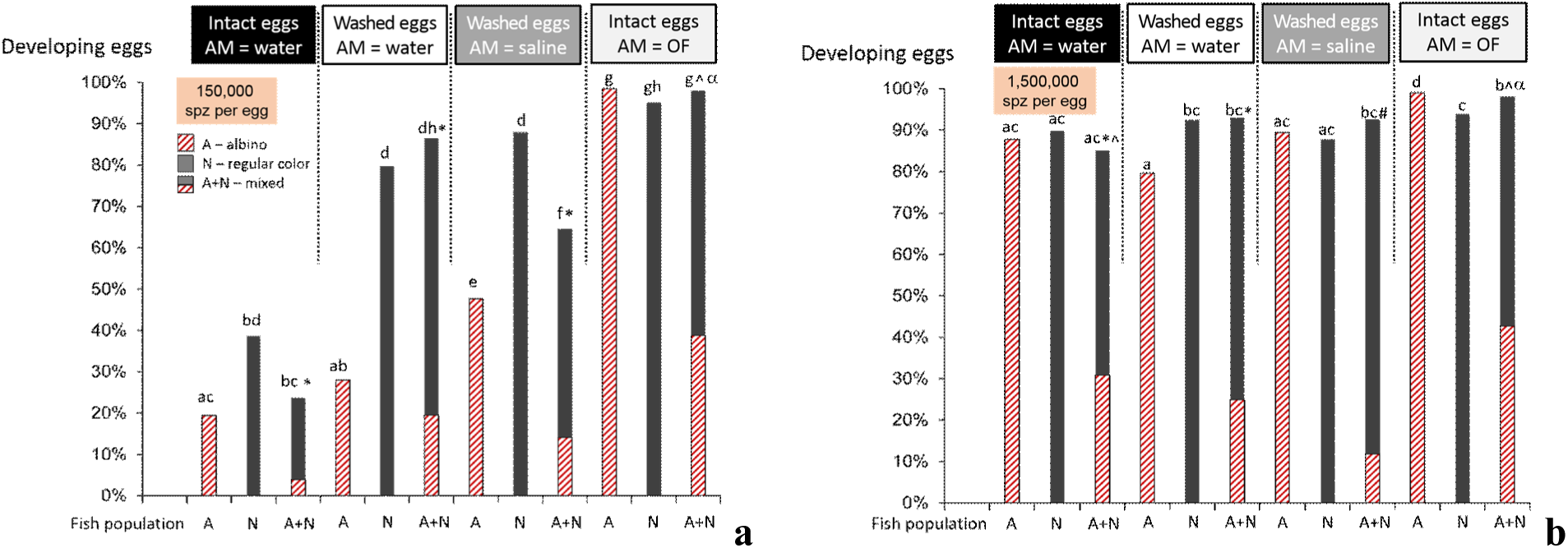
Ovarian fluid enhances the *in vitro* fertilization outcome. Albino rainbow trout egg embryo development rate after *in vitro* fertilization depends on the presence of ovarian fluid around the eggs or in the activation medium and male type (albino or regular color). Eggs (mixed from 3 females) were either non-washed to remove ovarian fluid (“Intact eggs”) or washed thrice with 0.9% NaCl solution (“Washed eggs”). Spermatozoa (a: 0.5 µl, ∼150,000 spermatozoa per egg in the activation medium, and b: 5 µl, ∼1,500,000 spermatozoa per egg in the activation medium) from 5 albino males (A), 5 regular color males (N), or the mixed sample (A+N) were activated in 8 ml of water, isotonic NaCl saline, or ovarian fluid (AM = water, saline or OF, correspondingly) and added to the beakers with eggs, mixed and the beakers were put to the shaker for 1 minute. After that, the eggs were rinsed, transferred to glass Petri dishes, and put to incubators at 11°C. In 11 days, the developing embryos were counted, and the fertilization rate was calculated (fertilized eggs/total amount of eggs), as well as the color type of the embryos was established. The striped bars show the albino embryos; plain color bars denote regular color embryos. Statistical significance of the differences was tested with the binomial model; due to multiple comparisons, the threshold p-value was set to 0.0005; different superscripts denote significant differences: Latin letters, between the overall development rates; signs * and ^, between albino embryo shares between the mixed groups; and α shows the absence of significant differences with the expected 50 to 50% share between albino and normal color embryos in the mixed groups.

### Ovarian fluid prolongs the motility of spermatozoa

It is widely accepted that kinetic characteristics of the spermatozoa, in particular their velocity, contribute significantly to the success of fertilization (*e.g.,* reported by Levitan [24]). It was mentioned above that ovarian fluid was found to improve the velocity of spermatozoa in several salmonid species [20].

Our results confirmed the significant effect of ovarian fluid on the velocity of rainbow trout spermatozoa (supplementary Figure S1), nevertheless not in terms of its increase in the initial motility period. In these experiments, rainbow trout spermatozoa were fully activated either in hypotonic media (water) or in isotonic media (ovarian fluid or physiological solution). The presence of ovarian fluid improved the longevity of spermatozoa, and the changes in the velocity of the spermatozoa activated in isotonic saline differed from both above media. To characterize these differences numerically, we performed a regression analysis of the experimental data.

The obtained linear regressions of average velocity changes during the motility period in all experimental conditions were characterized by high R^2^ (min 0.8, max 0.95; mean 0.88, SD 0.04, n = 17). Thus, these lines may characterize velocity changes during the post-activation time (Figure 2), and intercepts with the axes may serve as characteristics to compare spermatozoa’s initial velocity and longevity in different treatments. The indices characterizing the linear regressions are shown in supplementary Table S1, *i.e.*, the coefficient of determination (R^2^), the slopes, and the intercepts with the x (time) and y (velocity) axes. According to these data, the highest initial velocity was observed in the isotonic NaCl saline, while the lowest one was observed in the ovarian fluid. The period of sperm motility was most down in water and highest in ovarian fluid: the estimated longevity was almost 47, 67, and 74 s in water, isotonic saline, and the ovarian fluid correspondingly (Table S1).

**Figure 2.**
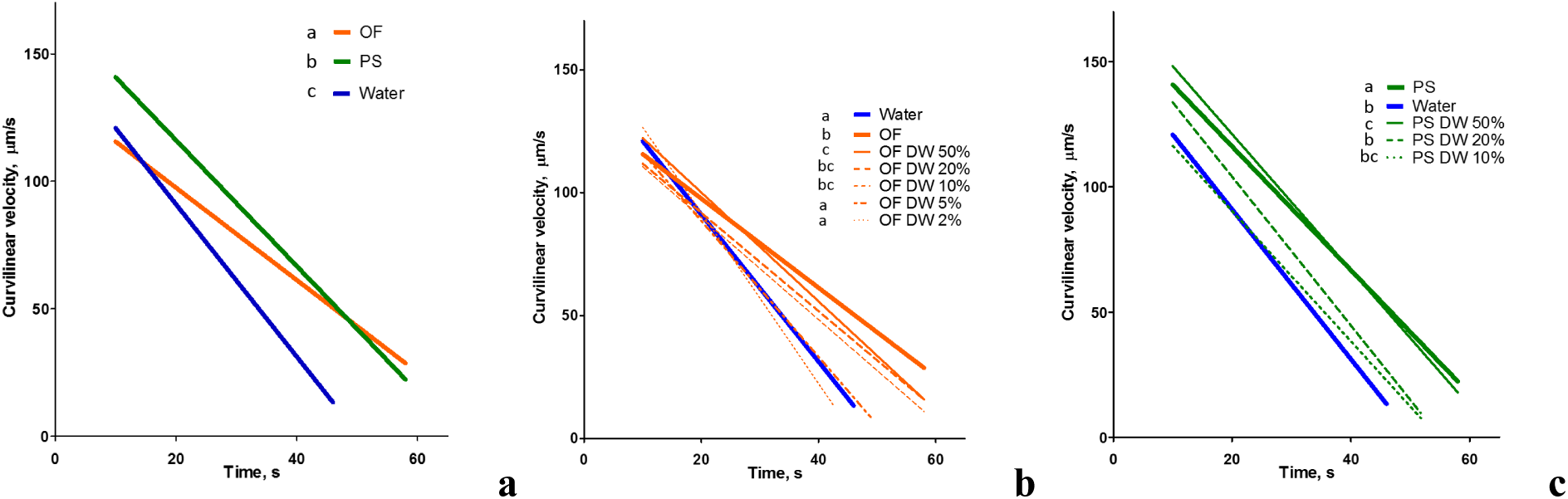
Ovarian fluid prolongs the motility of rainbow trout spermatozoa. The graphs represent changes in curvilinear velocity of spermatozoa activated in the presence of ovarian fluid and other conditions, depending on time post-activation: (a) velocity recorded in water, ovarian fluid, NaCl solution isotonic to ovarian fluid (physiological solution, PS, 290 mOsm/l); and dilutions of OF (b) and PS (c) with water. Data are linear regressions of experimental dependencies (shown in the supplementary Figure S1), the parameters of fitting, values of slopes and intercepts are shown in Table S1. Different superscripts denote significant differences between the slopes of regression lines in one graph, tested with ANCOVA and corrected for multiple comparisons, p<0.001.

**Table 1.**
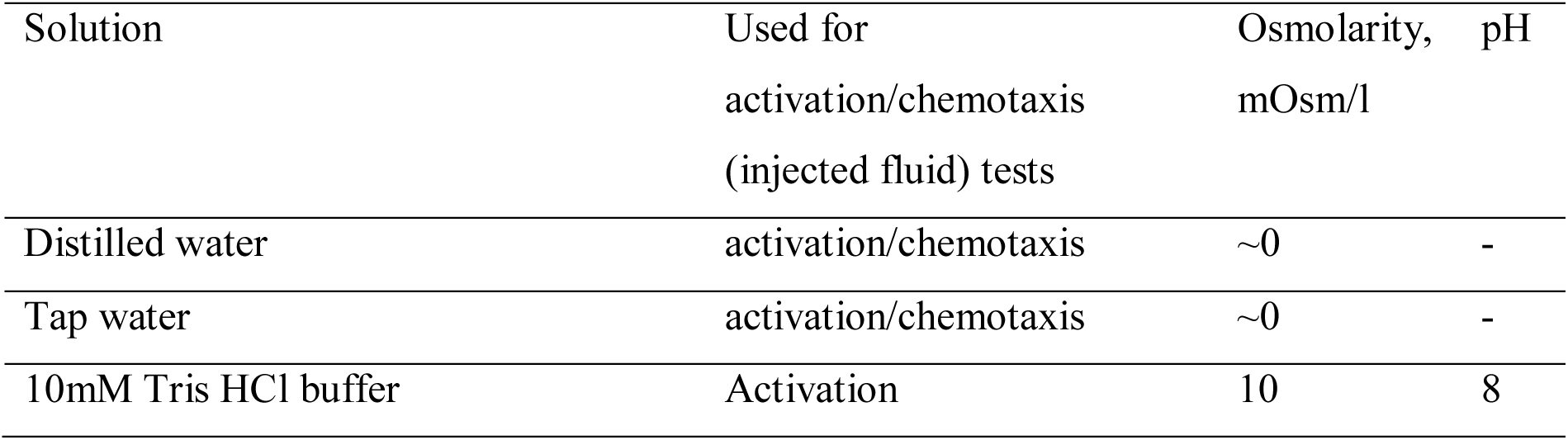
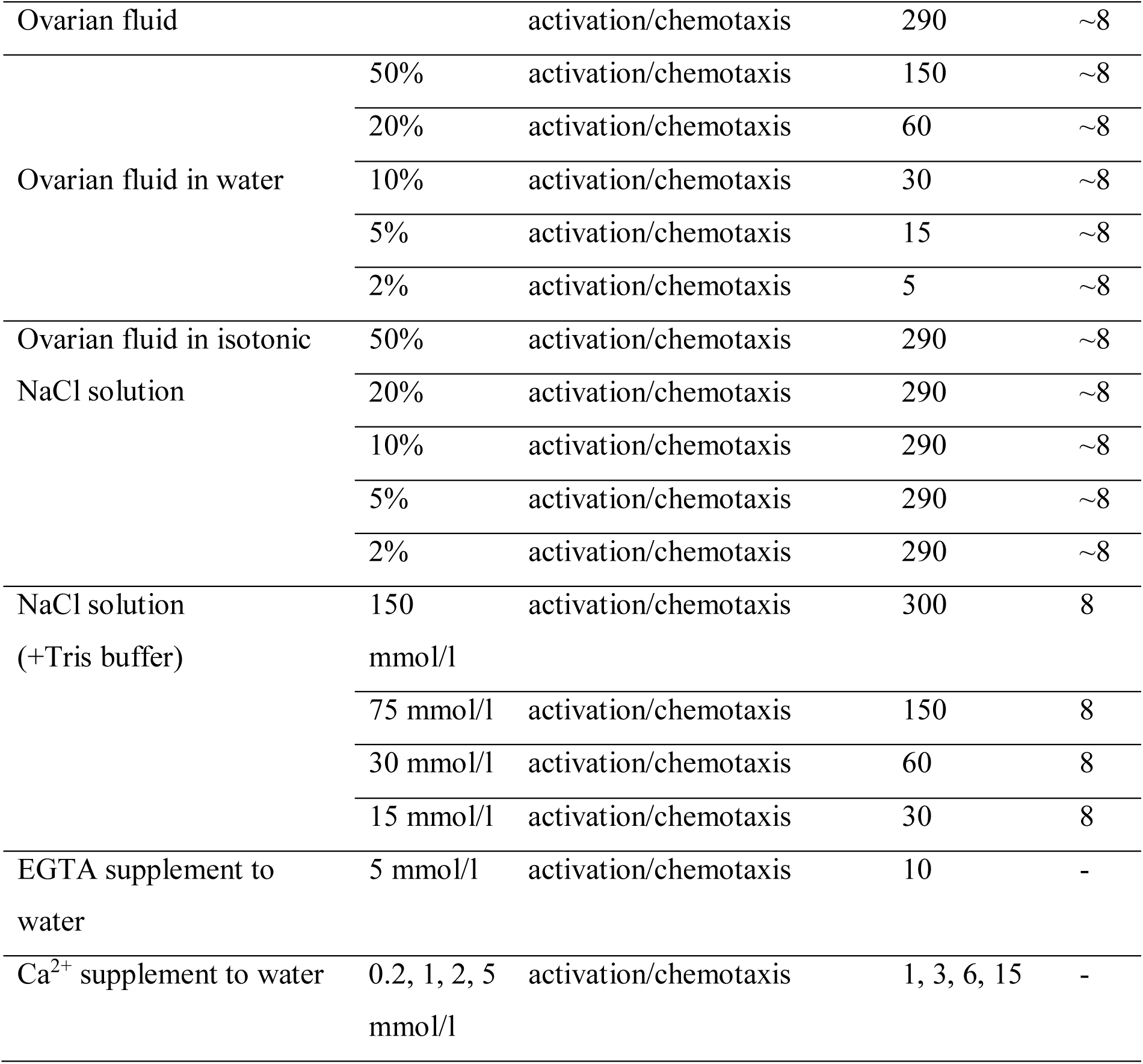
Solutions used for rainbow trout spermatozoa activation and/or chemotaxis tests

| Solution | Used for<br>activation/chemotaxis<br>(injected fluid) tests | Osmolarity,<br>mOsm/l | pH |
| --- | --- | --- | --- |
| Distilled water | activation/chemotaxis | ~0 | - |
| Tap water | activation/chemotaxis | ~0 | - |
| 10mM Tris HCl buffer | Activation | 10 | 8 |

|  |  |  |  |  |
| --- | --- | --- | --- | --- |
| Ovarian fluid |  | activation/chemotaxis | 290 | ~8 |
| Ovarian fluid in water | 50% | activation/chemotaxis | 150 | ~8 |
|  | 20% | activation/chemotaxis | 60 | ~8 |
|  | 10% | activation/chemotaxis | 30 | ~8 |
|  | 5% | activation/chemotaxis | 15 | ~8 |
|  | 2% | activation/chemotaxis | 5 | ~8 |
| Ovarian fluid in isotonic NaCl solution | 50% | activation/chemotaxis | 290 | ~8 |
|  | 20% | activation/chemotaxis | 290 | ~8 |
|  | 10% | activation/chemotaxis | 290 | ~8 |
|  | 5% | activation/chemotaxis | 290 | ~8 |
|  | 2% | activation/chemotaxis | 290 | ~8 |
| NaCl solution (+Tris buffer) | 150 mmol/l | activation/chemotaxis | 300 | 8 |
|  | 75 mmol/l | activation/chemotaxis | 150 | 8 |
|  | 30 mmol/l | activation/chemotaxis | 60 | 8 |
|  | 15 mmol/l | activation/chemotaxis | 30 | 8 |
| EGTA supplement to water | 5 mmol/l | activation/chemotaxis | 10 | - |
| Ca<sup>2+</sup> supplement to water | 0.2, 1, 2, 5 mmol/l | activation/chemotaxis | 1, 3, 6, 15 | - |

The dilution of ovarian fluid with water led to the disappearance of its longevity enhancing effect: if the content of ovarian fluid was 5%, there was no significant difference (p=0.88) compared with water in terms of regression line slope (Figure 2b, Table S1). If the spermatozoa were activated in the salines with the descending osmolarity, the resulting velocity changes were different from the dilutions of ovarian fluid (Fig 2c, Table S1).

### Ovarian fluid straightens the trajectories and has a trapping effect on the rainbow trout spermatozoa

The characteristics of spermatozoon trajectories, *e.g.*, the direction of their motion, are no less critical for the proper fulfillment of the male gametes’ function. It is evident from our findings, that rainbow trout spermatozoa are highly sensitive to the type of activating medium. Spermatozoa activated in water move according to tight circles during the initial period of motility (Figure3a). Trajectories of most spermatozoa swimming in ovarian fluid and NaCl isotonic solution are straight or arc-like with a larger proportion of straight moving cells in the ovarian fluid, which is reflected in a higher initial average linearity on the corresponding graph in Figure 3b. A two-fold dilution of ovarian fluid and NaCl isotonic solution did not change the path linearity essentially, while further dilution of both activation media dramatically decreased the proportion of spermatozoa moving straightforwardly. Interestingly, the pattern of motility in the diluted ovarian fluid has specific features compared to the saline medium with the same osmolarity: in the ovarian fluid, many cells move in complex coil-like trajectories (arrowheads in Figure 3a). These trajectories have the features of the so-called ‘turn-and-run’ pattern [25], which is associated with ‘explorative behavior’ of spermatozoa resulting from the effect of a chemotactic agent. There were no such explorative movement patterns observed in NaCl solutions with similar osmolarities.

**Figure 3.**
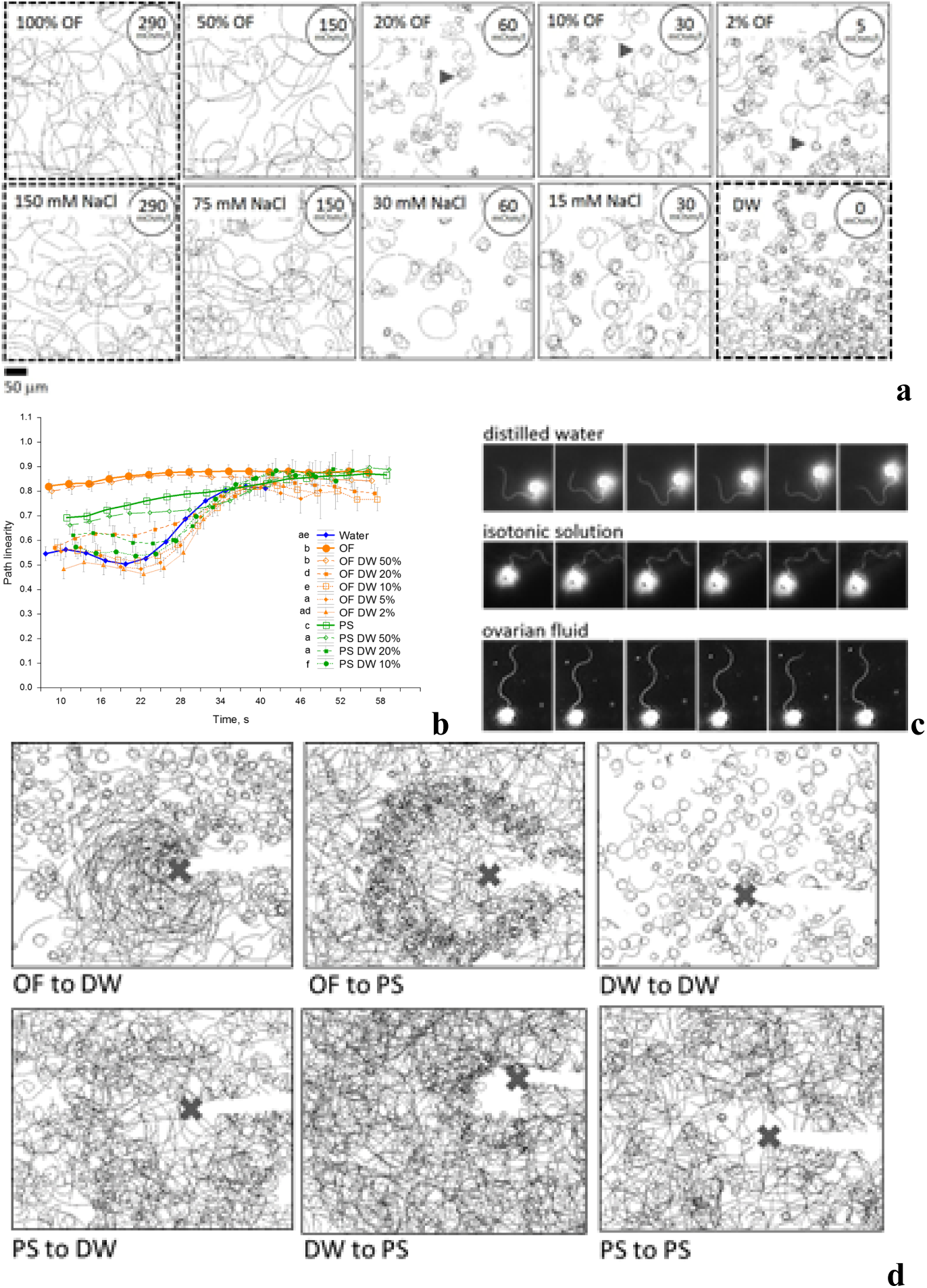
Ovarian fluid straightens the trajectories and has a trapping effect on the rainbow trout spermatozoa. (a) Spermatozoon trajectories visualized from 10 to 12 seconds post-activation in distilled water (DW), ovarian fluid (OF), NaCl solution isotonic to ovarian fluid (290 mOsm/l); and their dilutions with water (50; 20; 10; 5 and 2%). (b) The graph shows the linearity of swimming paths of rainbow trout spermatozoa activated in water, ovarian fluid, isotonic saline, and OF dilutions with water (50; 20; 10; 5 and 2%) depending on time post-activation. Data are mean values (n=16 for OF and 12 for PS dilutions; n = 26 for OF, PS and water), vertical bars denote 0.95 confidence interval; different letters in the legend denote significant differences, p<0.001. (c) The successive frames represent flagellar beating in rainbow trout spermatozoa activated in distilled water, 0.9% NaCl solution, or ovarian fluid. The frames are extracted every 0.005 s from the sequence recorded by high-speed video microscopy (at 2000 fps) using a 20× lens and darkfield microscopy. (d) Chemotactic sperm accumulation test: the frames show the swimming tracks of rainbow trout spermatozoa activated in various media near the tip of a microcapillary filled with test fluids: ovarian fluid (OF); distilled water (DW); isotonic NaCl saline (PS, 290 mOsm/l). Each track represents 2 seconds of motility (15-17 s post activation). The cross shows the tip of the microcapillary.

The flagellum is the only structure of the spermatozoon that may drive it and control the direction of its motility. The flagellum of motile rainbow trout spermatozoa has a typical wave shape (Figure 3c). There are differences in the distribution of the bends in the flagella of spermatozoa activated in water, isotonic saline, and ovarian fluid. The spermatozoa in the ovarian fluid have symmetric bends along the flagella, propagating uniformly. The flagella of the spermatozoa activated in isotonic NaCl saline have slight asymmetry of the flagellar waves, and this asymmetry is maximal in the cells activated in water. These different patterns are concordant with various modes of propagation of the cells in multiple media, *i.e.*, straightforward in ovarian fluid, straight to arc-like motion in isotonic saline, and tumbling in distilled water.

The behavior of spermatozoa in the media with spatially variating substances (gradients) may differ from the case with their homogenous distribution in the activation medium like in the above conventional motility tests. Such conditions may be created in the so-called spermatozoon accumulation test with a microcapillary, *i.e.*, using a glass capillary to introduce various test solutions into the bulk volume of medium containing activated spermatozoa.

Typical motility patterns of spermatozoa in our microcapillary tests are presented in Figure 3d and supplementary Video S1. No changes in the usual tumbling behavior of spermatozoa activated in water were observed when the microcapillary injected the same water. If the capillary was filled with ovarian fluid, the male gametes, which got into contact with the injected “cloud”, changed their behavior to a straight-line pattern. Moreover, the affected cells abruptly changed the direction of motion when reaching the border of the cloud, *i.e.,* they were trapped in the ovarian fluid, and a “positive taxis” was observed. Injection of ovarian fluid into the suspension of spermatozoa activated in isotonic NaCl resulted in the appearance of the “tumbling layer” of spermatozoa around the cloud and the part of the cells which entered the cloud moved straight. Changes in the behavior of spermatozoa and trapping were observed as well if isotonic saline was injected into the water as an activation medium. Nevertheless, no differences were seen if the activation media was changed to the same NaCl isotonic saline, which may suggest the presence of a sort of “osmotaxis” in our experimental conditions. Interestingly, injection of distilled water into the sperm suspension activated in isotonic saline caused the “negative taxis”: the cells avoided entering the hypoosmotic area, tended to stay in the medium with higher osmolarity, and performing abrupt turns on the borders of the “cloud” with water.

### Rainbow trout spermatozoa react with abrupt Ca^2+^ rise in flagella during turn-and-run behavior

Changes in the internal concentration of ions, Na^+^, K^+^, Ca^2+,^ and H^+^ are considered the critical factors of membrane hyperpolarization control due to osmotic pressure changes [26–28]. The presence of Ca^2+^ ions inside the spermatozoa was shown as an indispensable condition for motility initiation in most fish species [29]. Ca^2+^ takes part in the complex membrane cascade controlling the motility of spermatozoa and, what is most important, its direction [30, 31]. It was shown that the chemotactic response of ascidian spermatozoa, *i.e.,* the appearance of a specific “turn-and-run” pattern, similar to that observed by us, is associated with a burst-like rise in Ca^2+^ concentration in the bent segment of the spermatozoa flagellum [32].

Series of experiments similar to the above-described microcapillary tests were performed, which involved fluorescent microscopy to observe the changes in intracellular Ca^2+^ concentration after Fluo4 dye loading and further observation by fluorescence microscopy by light excitation at 435 nm. Observation of cells that perform swirling moves did not show any changes in the Fluo-4 fluorescence intensity of their flagella during motility (Figure 4a). In contrast, cells which exhibit a straight run and turn on the edge between the different media, *e.g.*, water and ovarian fluid or water and isotonic saline, show a bright “burst” of Fluo-4 fluorescence in the bent area of their flagella when turning (Figure 4b), *i.e.*, there was a local increase in Ca^2+^ concentration during the change of motility direction, associated with the appearance of asymmetric flagellar waves.

**Figure 4.**
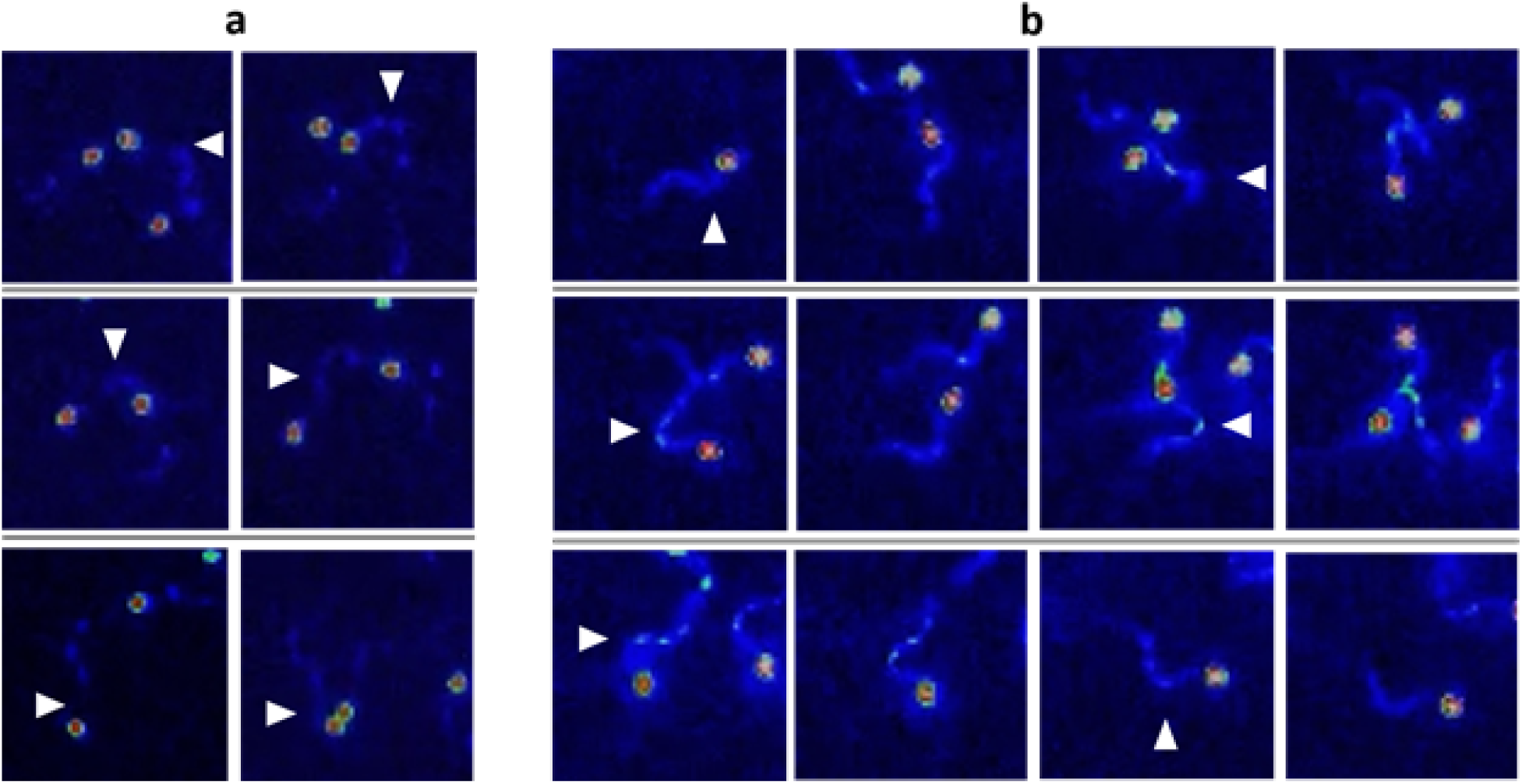
Rainbow trout spermatozoa react with abrupt Ca^2+^ concentration rise in flagella during ‘turn-and-run’ behavior. Fluo-4 fluorescence in t he sperm activated in water in the vicinity of the microcapillary filled with ovarian fluid or other fluids. Heat map: blue color – minimal concentration, red color – maximum concentration of Ca^2+^. (a) Frame by frame sequence of cell performing tumbling (shown by arrowheads). (b) - Frame by frame sequence of cell performing straight run and turns (shown by arrowheads).

### Isolated agents of ovarian fluid differently affect the behavior of rainbow trout spermatozoa. Effect of ovarian fluid molecular weight fractions and other protein solutions

It was reported in several species that maternal fluids may contain specific substances, which affect the spermatozoa, in particular, their chemotactic response [33].

What may be the active agent of the effect of ovarian fluid on the rainbow trout spermatozoa in addition to the mentioned above osmotic gradients? To answer this question, we have checked the performance of the male gametes in several molecular weight cut-off (MWCO) fractions separated from the ovarian fluid, as well as some other protein-containing fluids, *i.e*. thermo-treated ovarian fluid, rainbow trout blood serum, and an aqueous solution of bovine serum albumin.

Among the molecular weight fractions, only the 100+ kDa fraction caused the same motility and path linearity pattern as ovarian fluid (Figure5a, b). The traits in the other MW fractions (0-3, 3-10, 10-30, 30-50, and 50-100 kDa) did not differ significantly from the isotonic NaCl solution. The same motility pattern as in ovarian fluid were found both in rainbow trout blood serum and in the thermally treated ovarian fluid. In the aqueous solution of 1 mmol/l bovine serum albumin, the spermatozoa had the same trajectories as in water.

Regarding linear regression slopes, the molecular weight fractions of ovarian fluid were similar to NaCl isosmotic medium, except 0-3 and 100+ kDa fraction, and differed from OF (Figure5b).

In the microcapillary cell accumulation test, the low molecular OF fraction caused a quite bright response in the behavior of spermatozoa activated in water: some cells swirled in the vicinity of the border of the injected cloud, some spermatozoa moved straight without “escaping” the area, and some cells gathered close to the microcapillary tip (Figure 5c). Other fractions did not differ significantly from the isotonic saline in terms of the behavior of spermatozoa. Surprisingly, the introduction of blood serum caused the inhibition of the motility of cells entering the injected cloud. Introduction of thermo-treated OF entailed bright reaction associated with significant visible “border area”, *i.e.* the area where the cells were trapped and moved close to the source of the attractant.

**Figure 5.**
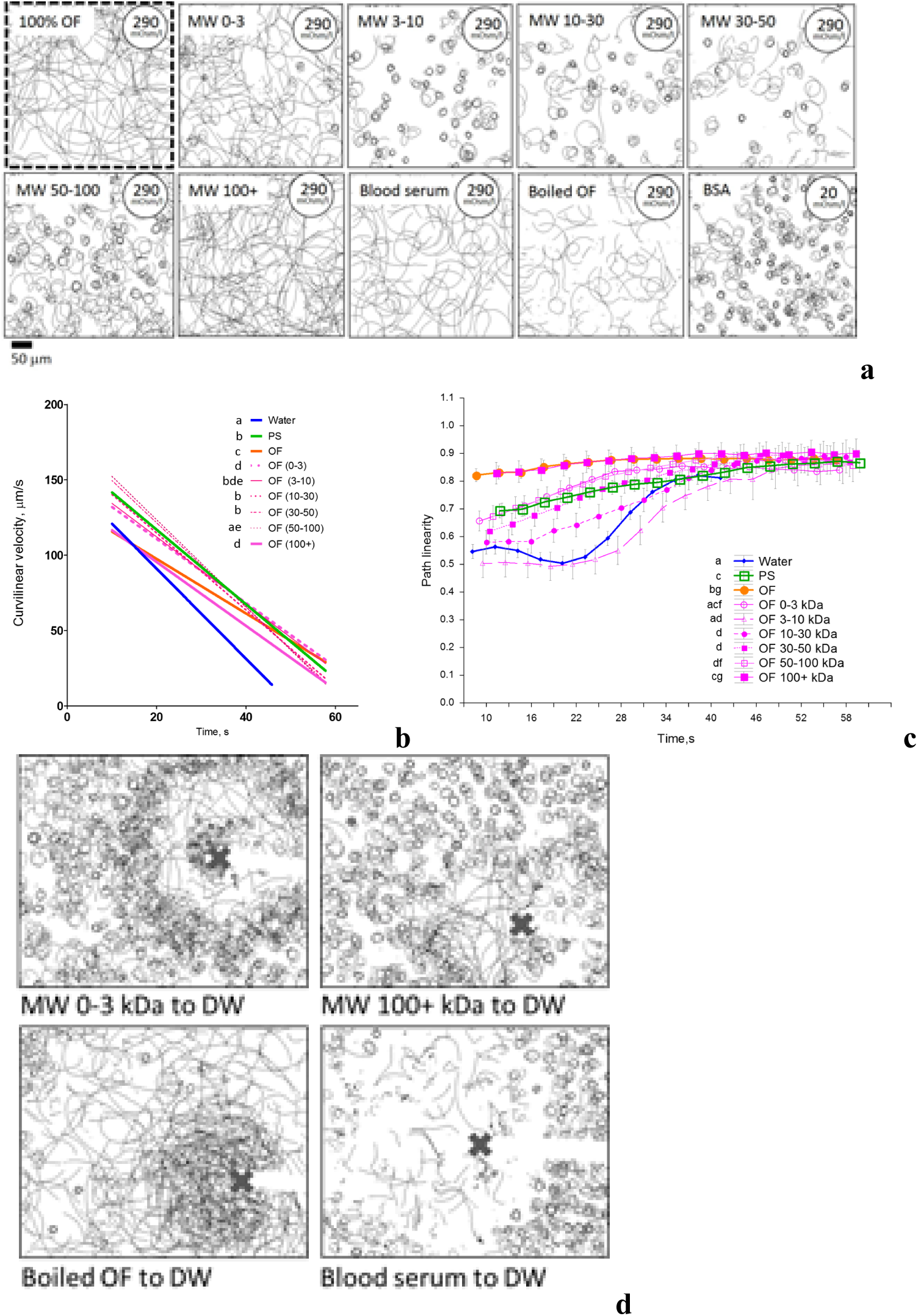
Isolated agents of ovarian fluid differently affect the behavior of rainbow trout spermatozoa. Effect of ovarian fluid molecular weight fractions, and other protein solutions. (a) Spermatozoon trajectories visualized from 10 to 12 seconds post-activation in MWCO fractions of ovarian fluid, blood serum, and thermotreated ovarian fluid. (b) and (c) The graphs show the changes in curvilinear velocity and path linearity of the spermatozoa during time post-activation in MWCO fractions of ovarian fluid, blood serum, and thermotreated ovarian fluid. Data are VCL linear regressions in (b) and path linearity mean values in (c); vertical bars denote 0.95 confidence interval; different letters in the legends indicate significant differences, p<0.001. Data are average for 26 males in water, ovarian fluid and NaCl isotonic saline, and 11 males in case of MWCO fractions of ovarian fluid. (d) Chemotactic sperm accumulation test: the frames show the swimming tracks of rainbow trout spermatozoa activated in distilled water near the tip of a microcapillary filled with test fluids: MWCO fractions of ovarian fluid with MW 0-3 kDa and 100+ kDa; blood serum and thermotreated ovarian fluid. Each track represents 2 seconds of motility (15-17 s post activation). The cross shows the tip of the microcapillary.

### Medium osmolarity and Ca^2+^ content have a “cross effect” on motility and trapping of rainbow trout spermatozoa

One of the suppositions about the role of ovarian fluid in the reproduction of salmonids is to provide a proper environment for the gametes’ interaction [16]. The osmolarity of the media is an important agent, triggering the motility onset and affecting its course, as shown above. Another essential agent involved in the motility phenomenon of fish spermatozoa is the contribution of calcium ions. The next series of experiments were done to assess the combined action of alternating external Ca^2+^ concentration and osmolarity on the motility traits of rainbow trout spermatozoa, and to compare these with the corresponding indices found in the case of ovarian fluid. Figure 6a shows the typical tracks recorded at 10-12 seconds post-activation of spermatozoa and combined with corresponding dependences of their curvilinear velocity in media supplemented by Ca^2+^ concentrations, varying from 0 to 5 mmol/l; and osmolarity of the media from 0 up to 300 mOsm/l. These indices include the most likely range of osmolarities and Ca^2+^ concentrations met by rainbow trout spermatozoa after their release by a male in natural conditions.

**Figure 6.**
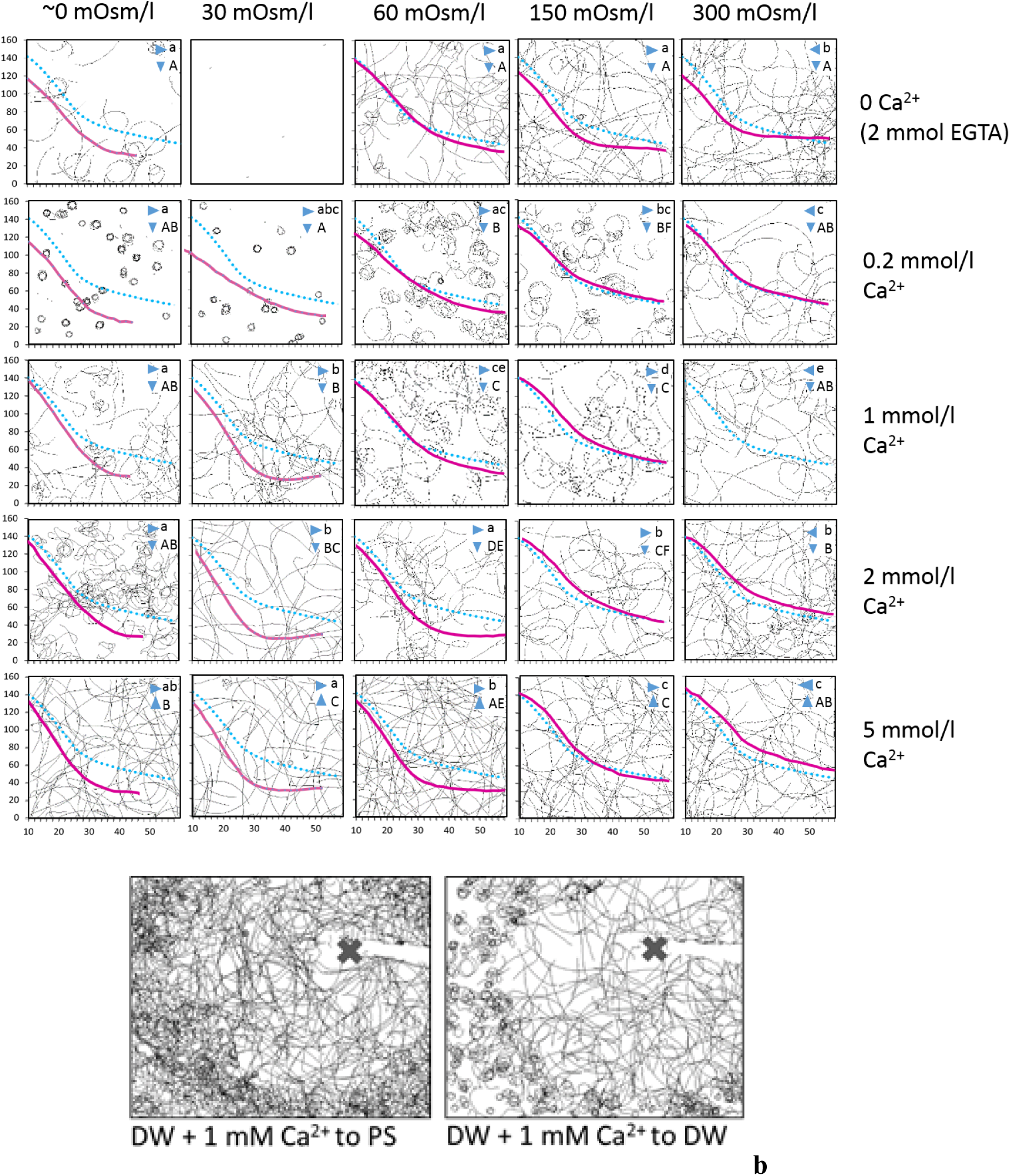
Medium osmolarity and Ca^2+^ content have a cross effect on motility and trapping of rainbow trout spermatozoa. (a) *Background*: spermatozoon trajectories visualized from 10 to 12 seconds post-activation in the media with osmolarities 30, 60, 150, 300 mOsm/l, prepared with NaCl and 10 mM Tris buffer, pH 8; and Ca^2+^ content 0, 0.2, 1, 2, and 5 mmol/l prepared with an appropriate amount of CaCl_2_ or 2 mmol/l EGTA in case of 0 mmol/l Ca medium. The “0” mOsm/l media was the distilled water with either 2 mmol/l EGTA or corresponding CaCl_2_ concentration, *i.e.*, the osmolarity in this media was, in practice, higher than 0; the actual values are shown in Table 1. *Curves*: solid lines show corres ponding curvilinear velocity changes of the spermatozoa *vs*. time post-activation in the described conditions. A dotted line is given for reference and shows the velocity recorded in the activation medium with 1 mmol Ca^2+^ and 300 mOsm/l osmolarity. Data are mean values (average from 8 males). The statistical significance of the differences between VCL dependencies was tested by pair-wise factorial repeated measure ANOVA with a threshold p-value of 0.0005. Different uppercase letters denote statistical differences in the media with varying Ca concentration at the same osmolarity (vertical), and different lower case letters indicate differences for media with varying osmolarity at the same calcium ion concentration (horizontal). (b) Chemotactic sperm accumulation test: the frames show the swimming tracks of rainbow trout spermatozoa activated in distilled water or isotonic saline near the tip of a microcapillary filled with distilled water and 1 mM Ca^2+^. Each track represents 2 seconds of motility (15-17 s post activation). The cross shows the tip of the microcapillary.

It is clear that the motility pattern depends both on the concentration of external Ca^2+^ and the osmolarity of the activation medium. In the absence of external Ca^2+^, the spermatozoa are poorly activated in 0 mOsm/l medium; the activated spermatozoa move in circular trajectories. Interestingly, the motility in the 30 mOsm/l medium was almost absent in all studied males. In the Ca-free media with 60, 150, and 300 mOsm/l osmolarities, the spermatozoa were activated, and most of them had straight trajectories. The curvilinear velocity of spermatozoa was similar in the beginning of the motility period; the cells in media with higher osmolarity had a longer period of motility and higher velocity after 30 seconds of motility period. The presence of 0.2 mmol/l Ca^2+^ dramatically changed the pattern of motility of spermatozoa. The majority of them moved according to tight circles in media with 0 and 30 mOsm/l osmolarity, and with a rise in osmolarity, the trajectories “uncoiled”, and become straighter in the 300 mOsm/l medium. In terms of curvilinear velocity, the cells in the media with 60, 150, and 300 mOsm/l osmolarities were faster and longer motile compared to 0 mOsm/l medium. The presence of 1 mmol/l Ca^2+^ in the activation media entails the straightening of the trajectories in most cases, even in the case of the 0 mOsm/l medium. Further rise of Ca^2+^ concentrations up to 2 and 5 mmol/l did not lead to significant changes compared to the case with 1 mmol/l concentration, except the medium with 60 mOsm/l osmolarity, where the VCL was surprisingly decreased faster, comparing to the media with lower Ca^2+^ content. The velocity graphs show the enhancing effect of media with 60, 150, and 300 mOsm/l osmolarities in terms of higher velocity and a more extended motility period in case of 1 mmol/l Ca^2+^, and 150 and 300 mOsm/l osmolarities in media supplemented by 2 and 5 mmol/l Ca^2+^.

The presence of Ca^2+^ ions also affects the response of the spermatozoa in the chemotactic microcapillary test, both in hypotonic and isotonic activation medium (Figure 6b). In the first case, the cells activated in the water change the motility pattern for a straight one after contact with the injected water containing 1 mmol/l Ca^2+^. The presence of calcium ions in the water injected to the spermatozoa activated in isotonic saline resulted in the absence of “negative osmotaxis”, mentioned above: the spermatozoa were able to enter the area with hypotonic medium and did not change the mode of the motility from the straightforward one.

## Discussion

The present study showed that ovarian fluid sustains chemokinetic effects on rainbow trout spermatozoa to affect their velocity, linearity, and longevity. The fluid exhibits a chemotaxis-like or trapping effect on sperm cells, and the male gametes show abrupt changes in the character and direction of motility due to the differences in the environmental conditions tending to follow more optimal conditions. These changes were accompanied by a burst-like rise in the Ca^2+^ content in the curved segment of the flagella. And finally, the ovarian fluid had a positive effect on the fertilizing ability of spermatozoa, improving the non-optimal motility of the cells. We will discuss how all these fit gametes’ encounter guidance in this species and first how the ovarian fluid presence changes the outcome of *in vitro* fertilization.

### Rainbow trout ovarian fluid provides an optimal environment for fertilization

The observation of a positive effect of ovarian fluid on the outcome of *in vitro* fertilization in salmonids was reported for Caspian brown trout, *S. trutta caspius* [34], and brown trout *S. trutta f. fario* [16]. On the contrary, Hugunin et al. [35] recommended wash-out the ovarian fluid before artificial fertilization in rainbow trout after finding the negative impact of ovarian fluid in “sub-fertile females”, which they explained as “preventing fertilization through impeding sperm movement or recognition/contact with the egg”. This adverse action might be associated with the substances released by overripen or destroyed eggs [36] on spermatozoa function. Several authors associated the discrepancies in the outcome of fertilization with a post-copulative effect rendered by females to select the best sire, or “cryptic female choice”. In particular, Butts *et al*. [37] have found significant enhancement of spermatozoon performance after activation in ovarian fluid from related females. Wojtczak *et al*. [38] have reported that the differential effect of ovarian fluid from various females is caused mainly by the inter-individual variation in pH of the maternal fluids; nevertheless, the authors mention that 40% of the variability is caused by other factors, which may include ions, carbohydrates or proteins. Rosengrave *et al*. [12] also found the differential effect of ovarian fluids from various females on spermatozoa performance in Chinook salmon *O. tshawytscha.* At the same time, there are opinions about the absence of cryptic female choice caused by ovarian fluid in the fertilization medium, *e.g.*, in Arctic charr reported by Kleppe *et al*. [39].

Our experiments have shown that using ovarian fluid as a fertilization medium significantly enhanced the fertilization outcome with sperm, which initially showed low motility in water, *i.e.* less than 20% motility in the albino male samples, and respectively low amount of fertilized eggs in case of 150.000 spermatozoa to 1 egg ratio (Figure 1a). This observation supports the opinions of Rosengrave *et al*. and Myers *et al*. [12, 20], which empha sized the inadequacy of pre-fertilization motility estimation in salmonids if it is performed in water.

Interestingly, in the case of non-washed eggs and water as the activation medium, the fertilization outcome for unrelated regular color males was the same low as for related albino males, despite much higher motility percentage of the former observed in water. Wash-out of ovarian fluid from albino female eggs significantly enhanced the number of regular color embryos (Fig 1a). If the ovarian fluid was used as the activation solution, the result of fertilization was maximal, as well as if isosmotic saline was used to activate the spermatozoa from both types of males. This probably shows that the performance of the spermatozoa may be triggered by the non-uniformity of the fertilization environment created by the layer of ovarian fluid confining the egg batch. Moreover, using ovarian fluid as a fertilization medium allowed to increase the percentage of embryos from related males, *i.e.,* the ovarian fluid increased the chances of spermatozoa from the males of the same population to win the sperm competition.

The above findings are valid for one of the two spermatozoon concentrations used in our study, *i.e.*, 150,000 spermatozoa per egg in the fertilization medium. Ten times rising of this ratio (to 1,500,000 spermatozoa per egg) entails the disappearance of most differences between experimental cases. This confirms the importance of adjusting the conditions of the *in vitro* fertilization experiments when performing the investigation of “tiny” associations and dependencies, which may be “masked” by an excess number of male gametes. Moreover, making associations with processes occurring in natural conditions may be inappropriate without exact knowledge of what are these natural conditions, *e.g.*, the ratio between gametes, *etc*.

Thus, the outcomes of *in vitro* fertilization experiments demonstrate the advantage of ovarian fluid in achieving the maximum fertilizing ability of spermatozoa. Below we consider some aspects of sperm behavior that potentially are related to understanding the mechanism of this phenomenon.

### Rainbow trout ovarian fluid enhances the kinetic traits of spermatozoa

Successful fertilization depends on the ability of the male gamete to carry the genetic material and fuse with the female gamete; thus, any change in the kinetic characteristics of the spermatozoa is highly essential. In agreement with the current opinion, one of the sperm characteristics, the spermatozoon velocity, contributes significantly to the success of fertilization. This was shown even in the case of a single male, *i.e.,* in the absence of competition between males, in this case, the outcome of fertilization correlated with a characteristic velocity of spermatozoa [40], and more clearly if the sperm competition was present, *i.e.,* the success of a particular male against rivals in terms of the bigger relative genetic contribution of the male with faster spermatozoa (*e.g*., reported by Levitan [24]).

Sperm competition is widespread in externally fertilizing vertebrate species, fishes in particular [41]. Salmonid fishes are typical examples of species with common and often intensive sperm competition. Their spawning tactics vary from spawning in pairs with the presence of parasitic or sneaker males to group spawning. The significance of spermatozoon velocity for the fertilization outcome in salmonids was reported widely (*e.g.*, by Gage *et al.*, and Liljedal *et al.* [42, 43]); in particular, Gage *et al.* [42] have found that relative sperm velocity is responsible for sperm competition success of a focal male compared to its rival and stated that the increased velocity of the “winner’s” spermatozoa allowed the faster search of the spawning microenvironment and penetration to the egg micropyle compared to the rival.

It was shown by several groups that the presence of ovarian fluid in the activation medium may positively affect the spermatozoa performance in these fishes by increasing the velocity of male gametes or their longevity [16,19,37,44–46]. Interestingly, Butts *et al*. [37] revealed in lake trout *Salvelinus namaycush* the rise in the curvilinear velocity of related (sibling) male spermatozoa after activating in 20% water solution of ovarian fluid compared to gametes from the unrelated male. However, Rosengrave *et al*. [46] have not found an y specificity driven by individual differences but revealed the absence of correlations between kinetic characteristics recorded in water and ovarian fluid.

In our study, we have confirmed that the presence of ovarian fluid affects the kinetic characteristics of the rainbow trout spermatozoa as compared to activation in water. Nevertheless, we have not found any rise of the curvilinear velocity at 10 s post activation, in contrast to the one reported by Turner and Montgomerie [45] in another salmonid, Arctic charr *Salvelinus alpinus*, and this was similar to the case of zebrafish, *Danio rerio*, reported by Poli *et al*. [19]. Like in the latter, the rate of rainbow trout spermatozoa velocity reduction during the post-activation period was much lower in the presence of ovarian fluid and its dilutions with water compared to plain water: significant difference disappeared only in case of 5% aqueous solution of ovarian fluid (Figure 2b, Table S1). The observed changes in the curvilinear velocity of spermatozoa activated in the isotonic saline differed significantly from that in the ovarian fluid. The initial velocity of the cells in saline was higher than both in water and ovarian fluid. Nevertheless, the longevity of spermatozoa activated in the ovarian fluid was the highest as compared to water or isotonic saline. This is in line with the “saving” effect of ovarian fluid reported by Elofsson *et al*. [47], which found a similar positive effect on sperm longevity in three-spined stickleback *Gasterosteus aculeatus* and associated it with solely t he ionic content of the fluid. Interestingly, the dilution of ovarian fluid with isotonic saline entailed the absence of significant differences in velocity between diluted (twice or more) ovarian fluid and the saline (Figure 2b).

In addition to the velocity of spermatozoa, the direction of their motility tracks plays an essential part in the propagation of the spermatozoa. The ovarian fluid had a prominent effect on the path linearity of the male gametes, which was highest compared to activation in water or isotonic saline. A similar observation about a significant rise in path linearity in rainbow trout spermatozoa activated in ovarian fluid compared to the saline buffer was reported by Dietrich et al. [17]. The straightening of the spermatozoon path in the ovarian fluid is associated with symmetrical propagation of the waves along the flagellum, unlike the non-symmetrical waves of the flagella in spermatozoa activated in water (Figure 3c). When decreasing the concentration of ovarian fluid, we observed a rise in the occurrence of specific trajectories, similar to the so-called “turn-and-run” pattern. The latter was frequently observed even in the highly diluted ovarian fluid, *i.e.*, 2% (Figure 3a) and in 5 and 10%, but was not present in non-diluted ovarian fluid or its 50% dilution. Such behavior was not characteristic of the sperm activated in isotonic saline.

Among the molecular weight cut-off fractions of the ovarian fluid, only the 100+ kDa fraction induced a motility pattern which presented no difference with that in the ovarian fluid, *i.e.*, the cells moved almost straightforwardly in both cases. In all other MWCO fractions, the trajectories were more circular, and the path linearity presented significant differences both with that of ovarian fluid and 100+ kDa fraction (Figure 5a, b).

### Ca^2+^ concentration and osmolarity have cross effects on motility traits

The osmolarity of the medium is one of the essential drivers of freshwater fish spermatozoon motility; in particular, the difference in osmolarity between outer and internal media initializes the function of the membrane orchestra of channels and other molecules and thus triggers the motility [48]. No less critical for motility initiation and its progress are the calcium ions, which are an integral part of numerous physiological processes occurring in the spermatozoon [49]. In our experiments on cross-effects of osmolarity and Ca^2+^ concentration, we varied these indices to identify which one among these isolated factors or their combination may be responsible for the chemokinetic effects exhibited by spermatozoa when contacting the ovarian fluid. It is clear that both factors (osmolarity and Ca^2+^ concentration) affect path linearity both individually and in combination, *e.g.,* in the absence of external Ca^2+^, the rise in the osmolarity of the activation medium from 0 to 300 mOsm/l entails the straightening of the spermatozoon trajectories, and rise in Ca^2+^ concentration from 0 to 5 mmol/l in hypoosmotic medium causes a similar effect. Remarkably, in the case of an intermediate concentration of Ca^2+^ (0.2 mmol/l), the path linearity was highly dependent on the osmolarity of the medium (Figure 6a). Collectively, the combination of Ca^2+^ concentration and osmolarity, which correspond to the ones in the ovarian fluid, controlled the motility traits similar to the effect of ovarian fluid, nevertheless the effect of solely osmolarity was the most significant.

### The ovarian fluid causes the attraction and trapping of spermatozoa

Microinjections of fluids into media containing activated spermatozoa is one of the obvious ways to mimic the interfaces occurring naturally between the water around the egg and the ovarian fluid that surrounds the latter. This allows modeling the behavior of male gametes during interaction with maternal fluids, which may differ from the motility characteristics expressed in a uniform environment. In our experiments, the rainbow trout spermatozoa were able to respond to the presence of this non-uniformity in the medium by becoming trapped within an area presenting more optimal conditions, *i.e.*, osmolarity, pH, and ionic content. In nature, these conditions may be provided only by ovarian fluid; nevertheless, in the experiment, it is possible to investigate how these different factors affect the response of the spermatozoa and which of them is/are the most critical.

Our experiments showed that positive taxis and trapping were observed towards the area with plain or diluted ovarian fluid for rainbow trout spermatozoa previously activated either in water or isosmotic saline. This effect was even stronger towards low molecular weight fractions or thermo-treated ovarian fluid, which is probably caused by an easier spatial distribution of active agents from the injected fluid with the viscosity lower than in ovarian fluid. It is interesting to note that in some cases, such as that shown in Figure 3d and 4c (and in supplementary Video S1), *i.e.,* for ovarian fluid injected to isotonic medium or low molecular fraction of ovarian fluid injected to water, spermatozoa separated into several populations, some of which showed a hyperactivation-like motility pattern close to the borders of the injected cloud or near the tip of the microcapillary. Changes of motility pattern for straightforward and trapping ones were observed as well in case of injection of isosmotic saline or some extent even for distilled water with 1 mmol/l Ca^2+^ (Figure 3d, 5c); these observations show that the overall positive taxis towards ovarian fluid results from a complex set of reactions of the rainbow trout spermatozoa to various factors, being the part of the ovarian fluid.

Interestingly, the rainbow trout spermatozoa were able to abruptly change the direction of their movement, *i.e.,* performing turn-and-run, when they met the interface between optimal and non-optimal environments. It may be exemplified by the trapping behavior in the case of injecting ovarian fluid or isotonic saline into the water with activated spermatozoa when the cells tended to stay inside the area with the conditions being optimal for motility (more extended period of straightforward motion). Moreover, a “negative taxis” behavior was observed in the case when distilled water was injected into the isosmotic activation medium: here, the male gametes “avoided” entering into the injected medium (Figure 3d). This behavior allows us to conclude that rainbow trout spermatozoa are highly sensitive to the environment and can change the direction of their motion to follow or to stay in the optimal conditions, which in nature are most likely provided by ovarian fluid.

### Effect of rainbow trout ovarian fluid on spermatozoa is in line with the effects of female factors in other externally fertilizing species

The phenomena accompanying the performance of rainbow trout spermatozoa in the presence of female fluids are reminiscent of previously described triggered motility of marine invertebrates, *e.g.,* sea urchins [50], ascidians [51], or squids [52]. The similarity of these sorts of behavior is even more spectacular considering that turn-and-run loops in rainbow trout spermatozoa were accompanied by asymmetric bending of the flagella and burst-like rise of Ca^2+^ concentration in the bent flagellar segment (Figure 4). A similar phenomenon was found to mediate the chemotactic activity of ascidian spermatozoa as its inherent part [10].

As was mentioned above, the vast majority of existing theories and models of spermatozoon chemotaxis are based on numerous investigations performed in marine invertebrates [9, 10]. Among the fishes, only the Pacific herring *Clupea pallasii* has relatively well-studied mechanisms of sperm-egg encounters so far. Yanagimachi *et al*. [6] have represented the prospective model of the herring spermatozoon motility initiation and entry into the egg. The spermatozoa may stay undamaged in the water prior to activation for several hours and even days, that is a unique feature of herring sperm among fishes [53]: the males first release their sperm into the water, and the females are attracted to the spot and release their eggs then. The male gametes are activated in the presence of a specific protein (defined as SMIF, sperm motility initiating factor) associated with the ovarian fluid and the egg membrane [54]. The activator molecule interacts with membrane receptors, which is followed by activation of adenyl cyclase, potassium and calcium ion channels, and respective accumulation of cAMP, influxes of K^+^ and Ca^2+^ [6]. After being activated, they are attracted to the micropyle area with the aid of another protein (MISA, micropylar sperm attractant) [55]. This attraction is accompanied by the effect of proton pumps and Ca^2+^ channels, which control the intracellular concentration of calcium ions and, correspondingly, the flagella beating pattern [6].

Unlike the Pacific herring, the rainbow trout spermatozoa can be activated in water; nevertheless, this activity is extremely short (as in most freshwater species). The male (or males) ejaculates in the exact moment or shortly after the female releases the eggs into the prepared nest together with plenty of ovarian fluid. The spermatozoa of rainbow trout change their motility pattern in the presence of ovarian fluid: the more ovarian fluid present in the activation medium, the straighter the path of spermatozoa. In the diluted ovarian fluid, the spermatozoa perform explorative “turn-and-runs”, and have positive taxis to the ovarian fluid. The cells, which entered these areas become trapped – if they reach the borders with water, they change the direction of motion, performing abrupt turns. This turn-and-run activity is accompanied by the appearance of asymmetric bends in the flagella and a flash-like increase of Ca^2+^ concentration in the bent segment of the flagellum, presumably accompanying the encounter of the spermatozoa towards the eggs. These phenomena have certain similarities to the ones described in the Atlantic herring, as well as in marine invertebrates, which allows us to suppose the common basis of chemotactic responses in the broader spectrum of externally fertilizing animals than it was established before. The found individual peculiarities of the egg (ovarian fluid) – sperm interaction in rainbow trout may be associated with specific features of the spawning process in this species.

Thus, in the frames of one investigation, we have performed the detailed testing of the features accompanying the encounters in one of the representatives of freshwater spawning fish (rainbow trout *Oncorhynchus mykiss*). These tests revealed the triggering effect of female fluids on the performance of male gametes, which conforms to the features of spawning behavior in these species. We have not found a single specific agent responsible for the changes in the motility of spermatozoa, rather supposed the combined effect of several factors. Nevertheless, finding the exact nature of the active agents and the responsive structures in the male gametes is the prospective goal of future studies.

The studying of mechanisms underlying the reproductive processes is an important part of evolutionary developmental biology. These mechanisms adapt to the specific conditions of the environment both in the course of evolution and even the lifetime of an individual. The adaptations entail the arising and modification of reproduction strategies and tactics. These are incredibly various in fishes due to the different ecological niches which they inhabit. The fact that most of the fishes have the external mode of reproduction makes them a good model for performing various experiments in the conditions close to *in vivo*, or with the controlled shift of *in vivo* conditions, especially in the case of freshwater species, since in addition to other factors their process of fertilization should be highly synchronous and chronically limited to several minutes. Thus, the broader inclusion of fishes into future studies of the gamete encounters will significantly improve the overall understanding of the evolutionary aspects of reproduction, its molecular mechanisms, and their effects on reproductive behavior and *vice versa*.

## Materials and methods

### Ethical statement

The experimental protocols of the study were approved by the Institutional Animal Care and Use Committee at the University of South Bohemia, following the legislation on the protection of animals against cruelty (Act no. 246/1992 Coll.). Manipulations with animals were performed according to the authorization for breeding and delivery of experimental animals (Reference number: 56665/2016-MZE-17214 17OZ19180/2016-17214, valid from October 4, 2016, for 5 years) and the authorization for the use of experimental animals (Reference number: 2293/2015-MZE-17214 16OZ22302/2014-17214, valid from January 22, 2015 for 5 years) issued to the Faculty of Fisheries and Protection of Waters, the University of South Bohemia by Ministry of Agriculture of the Czech Republic. The study was carried out in compliance with the ARRIVE guidelines where appropriate (https://arriveguidelines.org).

### Fish broodstock. Gametes and fluids collection

The experiments were performed with gametes of rainbow trout *Oncorhynchus mykiss* (2-3-year-old, 0.6-1 kg weight) in the spring and autumn seasons in 2017, 2018, and 2019. Altogether samples from 60 fish were used in this study. Eggs and sperm were obtained during the natural spawning period from strains that spawn in spring or autumn. Fish were obtained from a fish farm in Bušanovice, the Czech Republic, and Oshino Trout Hatchery, Yamanashi Prefectural Fisheries Technology Center, Oshino, Yamanashi, Japan (in autumn 2018). The male fish were selected randomly from the farm stock, transported in oxygenized water tanks, and kept indoor in the covered water tanks with fresh water and oxygen supply at 11°C. Sampling was done in all spermiating males once a week. Sperm was collected by stripping, stored during analysis on ice, and used for 6 hours. The eggs were collected from the batches prepared for artificial propagation at the fish farm during planned fertilization procedures. Ovarian fluid was drained off from the eggs with a sieve, centrifuged to remove debris, and stored at 4°C in plastic tubes or frozen at −80°C when long-term storage was required. Only ovarian fluid from non-overripen eggs was used in conventional motility analyses. The fertilization test presented in the paper was done in February 2019 using eggs of albino rainbow trout females and sperm from albino and normal color males; ovarian fluid from albino females was collected and stored separately.

### Preparation of media

The evaluation of the chemokinetic properties of the ovarian fluid was done in a series of activating media (Table 1): with varying concentrations of OF (from 100 to 2%); with various concentrations of NaCl (Sigma Aldrich, USA), to mimic the osmotic pressure of corresponding dilutions of ovarian fluid; with different molecular weight fractions of ovarian fluid; thermo-treated ovarian fluid (ovarian fluid was boiled at 100°C for 5 minutes); rainbow trout blood serum (collected from the blood centrifuged at 3000g for 5 minutes); bovine serum albumin (Sigma Aldrich, USA) 1 mg/ml solution in water. Molecular weight cut-off (MWCO) fractions of the ovarian fluid were prepared using Amicon Ultra centrifugal filters with 3, 10, 30, 50, and 100 kDa filters (Merck Millipore Ltd., Ireland). Stepwise centrifugation through the filters was done starting from 100 kDa. Processing was performed at 4°C till 10-time concentrating the fluid above the filter (10% of the initial volume). The volume of the collected fluids was recovered with isotonic NaCl (0.9% w/v) solution. The fluid passed through the filter was transferred to the next filter with lower MWCO, and the procedure was repeated until the lowest MW filter was used.

Additional experiments were performed by checking the cross-effect of Ca^2+^ concentration and osmolarity of the activation media. For doing this, the set of media was prepared with varying osmolarity (0, 30, 60, 150, 300 mOsm/l, done with NaCl and 10 mM Tris buffer (Sigma Aldrich), pH 8, where appropriate) and Ca^2+^ content (0, 0.2, 1, 2, and 5 mmol/l made with the appropriate amount of CaCl_2_ (Sigma Aldrich) or 2 mmol/l EGTA (Sigma Aldrich)). The “0” mOsm/l media was distilled water with either 2 mmol/l EGTA or corresponding CaCl_2_ concentration, *i.e.,* the osmolarity in this media was, in reality, higher than 0, the actual values are shown in Table 1.

Appropriate solutions were assessed for osmolarity, pH, protein, and ion content. Osmolarity was measured using a freezing point osmometer Osmomat 3000 (Gonotec GmbH, Germany) and presented in mOsm/l. Concentrations of sodium and potassium ions were measured by potentiometry using ion-selective electrodes (Bayer HealthCare, Tarrytown, NY, USA). Calcium ion concentration was measured by absorption photometry applying the o-cresolphthalein complexone method [56]. The ion concentration is expressed in mmol/l of the medium. Protein concentration was determined using the Pierce BCA Protein Assay kit (Thermo Scientific, USA) and shown in mg/ml. The measurements of the protein and ion contents were done in the range of standard calibration curves, appropriate to the used method.

### Motility observation and recording

Sperm suspensions (around 0,1 µl) were carefully mixed for 5 s with 40 µl of tested solutions, and motility (if present) was recorded for 1 min post-activation using digital camera ISAS (PROISER, Spain) or IDS (IDS Imaging Development System GmbH, Germany) set at 25 frames/s and a microscope (UB 200i, PROISER, Spain) using 20x lens and negative phase-contrast. The experiments were performed at 7°C (supported by the cooling stage (Semic, Poland)). Video records were analyzed to estimate spermatozoa motility traits using ImageJ software (U. S. National Institutes of Health, Bethesda, Maryland, USA) and the following plugins: CASA and CASA modified for multiple analyses [57, 58]. The analysis was performed if the decline in the percentage of motile cells did not reach 10%; and only moving cells with a curvilinear velocity higher than 15 ¼m/s were considered as motile. Values of spermatozoa velocity, the linearity of the spermatozoa trajectories, as well as the pattern of motility (1-2 sec tracks of individual spermatozoa in the recorded field) were obtained.

### Sperm chemotaxis tests

A series of experiments were done to assess the response of the spermatozoa to the injection of various test fluids (see Table 1) into activating media using glass microcapillaries, which represent an accumulation assay conventionally applied for simple spermatozoa chemotaxis analysis [59]. For this purpose, glass microcapillaries (G100, Narishige, Japan) were pooled (PC-100 puller, Narishige) to get microneedles with tips of ∼20 µm openings size, which were additionally cut by a microgrinder (EG-401, Narishige) to obtain uniform tip openings. The microcapillary was filled with test fluid and assembled with a microinjector (CellTram Vario, Eppendorf, Germany), then fixed on a holder (Narishige) and adjusted above a specimen glass on a microscope stage. The microinjector pressure was applied to ensure the slow discharge of the fluid. A drop of the activation medium (40 µl) was placed on the glass slide, spermatozoa were activated in the drop by gentle stirring, and the microneedle with discharging fluid was introduced in it right after activation of spermatozoa (∼1-2 s). The behavior of the spermatozoa in the vicinity of the tip of the microcapillary was observed directly under the microscope and video-recorded for 1 minute. The resulting records were then processed by CASA plugin for ImageJ to get the tracks of spermatozoa, and these patterns of motility were thereafter analyzed.

### High-speed imaging of spermatozoon flagellar beating

The process of a flagellar beating during the motility of spermatozoa in different media was studied using high-speed recording [60]. Motility of spermatozoa was recorded during 10-20 s post-activation in water, isotonic NaCl solution, or ovarian fluid using darkfield microscopy (Olympus BX50, Japan). The records were performed by a high-speed video camera (model i-SPEED TR, Olympus, Japan) with 2000 frames per second rate.

### Ca^2+^ concentration imaging

For imaging of intracellular calcium ion content ([Ca^2+^]i), semen was suspended in 4–5 volumes of immobilization solution, *i.e.,* artificial seminal fluid (Morisawa and Morisawa 1988) containing 0.05% Cremophor EL (Dojindo, Japan) and 20 μM Fluo-4 AM (Dojindo), and incubated for dye loading at 10°C during 2 hrs. Then the sperm was activated in an observation chamber, and a microinjector with test liquid was introduced. Fluorescence signals emitted by the swimming sperm were captured by a microscope Olympus IX71 (Japan) equipped with fluorescence illumination and a digital CCD camera QImaging Retiga-EXi (Adept Turnkey, Australia) according to the method described by Shiba *et al*. [32]. The experiments were done at 12°C. The videos were then analyzed, and the fluorescence of individual cells was tracked by the imaging application TI workbench [61]. The typical fluorescent response observed in spermatozoa during motility is represented in a series of heat map images, where the blue color denotes the minimum fluorescence of Fluo-4, and the red color shows the maximum fluorescence.

### In vitro fertilization

The *in vitro* fertilization assays were performed in February 2019. The fertilization was done with eggs collected from 3 albino females and two mixed sperm specimens from 5 albino males or 5 normally colored males. The pooling of eggs from various females (as well as their ovarian fluids) was made to prevent individual male-female interactions, which was not the issue of this experiment [62]. The fish were selected randomly, as stated above. The pooling of eggs from various females is made to minimize the effects of individual male-female interactions [62], which was not the issue of this experiment. The experimental groups are shown in Table 2. The experimental design was aimed to check if the presence of ovarian fluid may change the outcome of *in vitro* fertilization. In all cases, 5 g of eggs (on average 79 eggs) were fertilized by 0.5 µl or 5 µl of sperm mixed in 8 ml of water from the hatchery supply system at 11°C. The concentration of spermatozoa in the sperm was 2.29×10^10^/ml, *i.e.*, around 150,000 and 1,500,000 spermatozoa per egg respectively for 0.5 and 5 µl sperm in the fertilization medium. In some cases, the eggs were washed with 0.9% NaCl solution three times during 10 sec to remove the ovarian fluid. The eggs (washed/non-washed) were put into the plastic beaker, poured with test solution (water, isotonic NaCl solution, or ovarian fluid) and the sperm was added. The beakers were then placed on a shaker (around 100 rpm) and after 1 minute-incubation the eggs were rinsed and transferred to glass Petri dishes. Each point was done in three technical replicates.

**Table 2.**
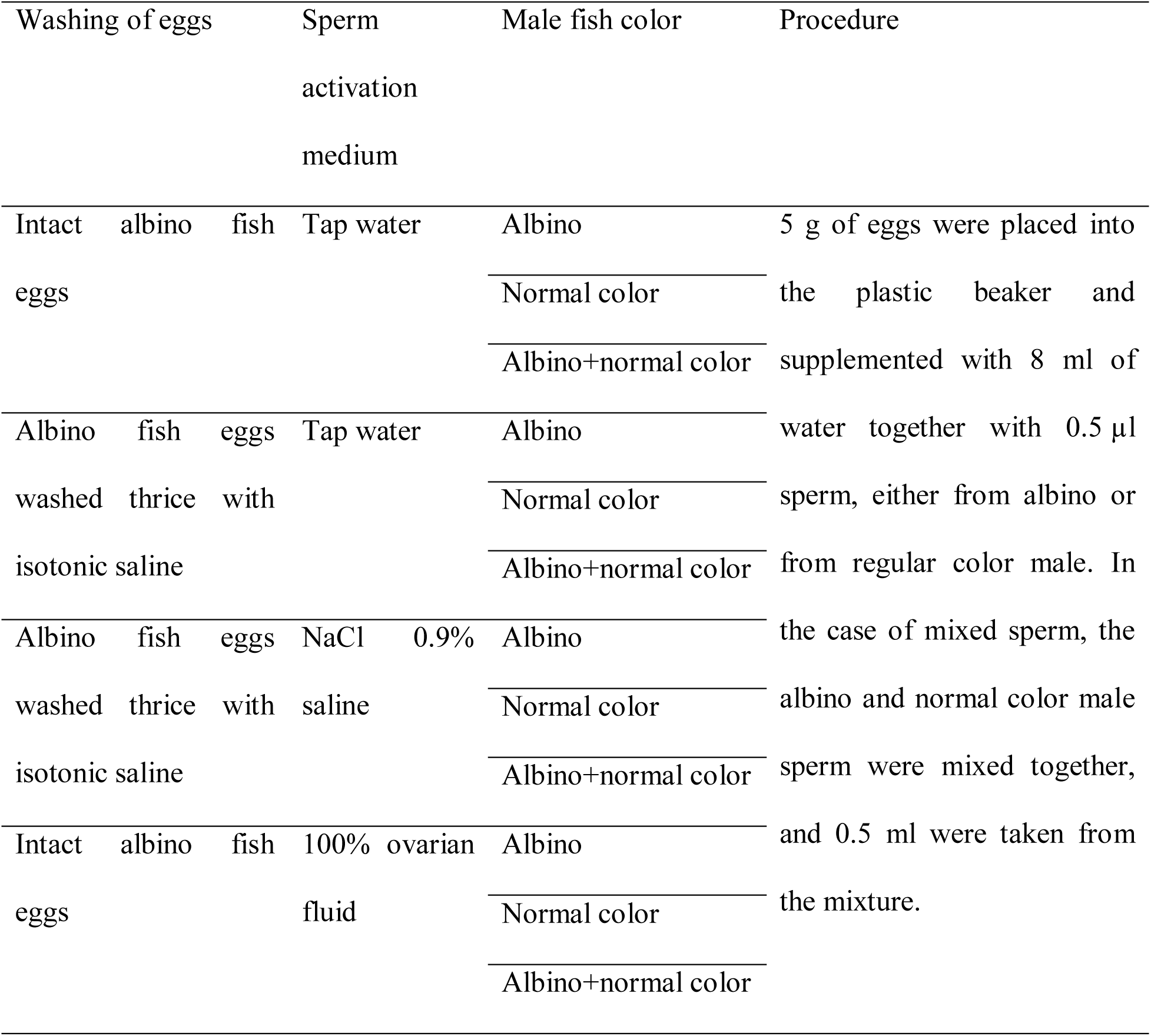
Effect of ovarian fluid on fertilization performance in rainbow trout, albino vs. conventional color fish: experimental design

The dishes were transferred into the tank with baskets for further incubation at 11°C. The tank had a closed water circuit with aeration, UV-treatment, and temperature control. All fertilizations were done in three replicates. The outcome of *in vitro* fertilization was assessed in 11 days by the amount of developing embryos (the embryo development rate is the amount of developing embryos divided by the total amount of eggs); simultaneously, the share of albino embryos was counted. The number of the hatched larva was counted as well. The hatching rate (number of hatched larva/number of eggs) did not differ significantly from the number of developing embryos. Thus, only the latter is presented.

### Statistical analysis

Assessment of motility parameters in different activation media was conducted in triplicates for 26 males in case of water, ovarian fluid, and NaCl isotonic saline (performed as a control during motility and chemotactic tests); 16 males in case of OF dilutions, 12 males in media with various osmolarity based on sodium chloride; and 11 males in case of MWCO fractions of ovarian fluid. Sperm samples from 8 males were used for experiments with combined effects of Ca^2+^ concentrations and osmolarity. Curvilinear velocity (VCL) and path linearity (LIN) were obtained from the motility measurement from 50-300 spermatozoa per replicate per time point during 10-59 seconds post-activation with 3 s increment. The motility parameters (average from technical replicates) were then *loge* and *logit* transformed (VCL and LIN, respectively) to ensure the normal distribution of the data, and analysis of interactive effects between variables was performed using repeated measures ANOVA in Statistica v. 13 (TIBCO Software Inc., USA). Media and post-activation time were considered as within-subjects factors and VCL or LIN as dependent ones. In the case of calcium ions/osmolarity cross effect, both calcium concentration and medium osmolarity and time were considered as within-subject factors. Pair-wise analysis was done to establish significant differences in interaction between media and time between various activation conditions (*i.e.*, the difference in spermatozoa behavior in multiple media along motility time was present). The corresponding correction of p-value was done to consider the multiple comparisons; the threshold p-value was set to 0.0005.

The data for spermatozoa velocities in several media (water, isotonic NaCl saline, ovarian fluid and dilutions in water, dilutions of isotonic saline with water, MWCO fraction of ovarian fluid) were used then to obtain linear regression dependencies in GraphPad Prism version 5 for Windows software (La Jolla, CA, USA); and the following parameters were obtained: slope (A), intercepts with x and y axes (B and C), coefficient of determination or the goodness-of-fit (R2). The hypothesis for the equality of regression slopes was checked using ANCOVA, and the threshold p-value was set to 0.0001.

Binomial model was used to assess significant differences between the groups in the fertilization test. Three technical replicates for each point were pooled together to increase the statistical power of the analysis. The analysis was done using G*Power 3.1.9.6. software [63]. Following hypotheses were tested: 1) if the obtained developing embryo proportions are the same among the groups; 2) if the proportion of albino embryos is the same between mixed groups; and 3) if the proportions of albino and normal color embryos are the same in the mixed groups. Considering the multiple comparisons, the threshold p-value was set to 0.0005.

## Acknowledgments

The study was financially supported by the Ministry of Education, Youth and Sports of the Czech Republic projects - „CENAKVA“ (LM2018099) and “Biodiversity” (CZ.02.1.01./0.0/0.0/16_025/0007370), by the Grant Agency of the University of South Bohemia in Ceske Budejovice (125/2016/Z, 013/2018/Z, and 097/2019/Z) and by the Czech Science Foundation (18-12465Y).

## Author contributions

V.K., J.C. and S.B. designed the research; V.K., B.D., M.R., M.Y., J.C., and S.B. developed the methodology; V.K., B.D., M.R., and S.B. conducted the experiments; V.K., B.D., and S.B. analyzed the data; V.K., B.D., H.G., M.Y., J.C., and S.B. wrote the paper.

## Annex

**Table S(supplementary)1.**
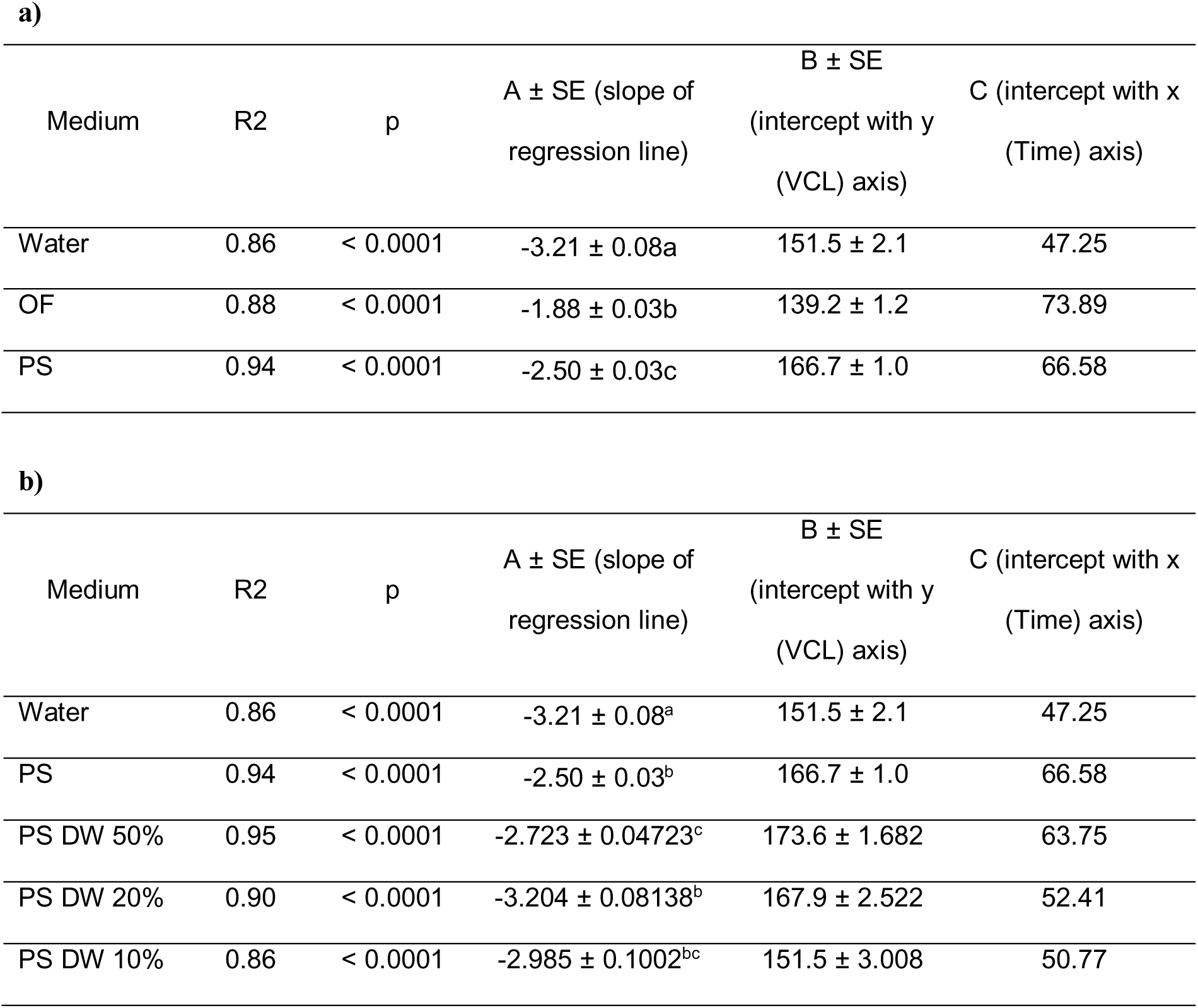
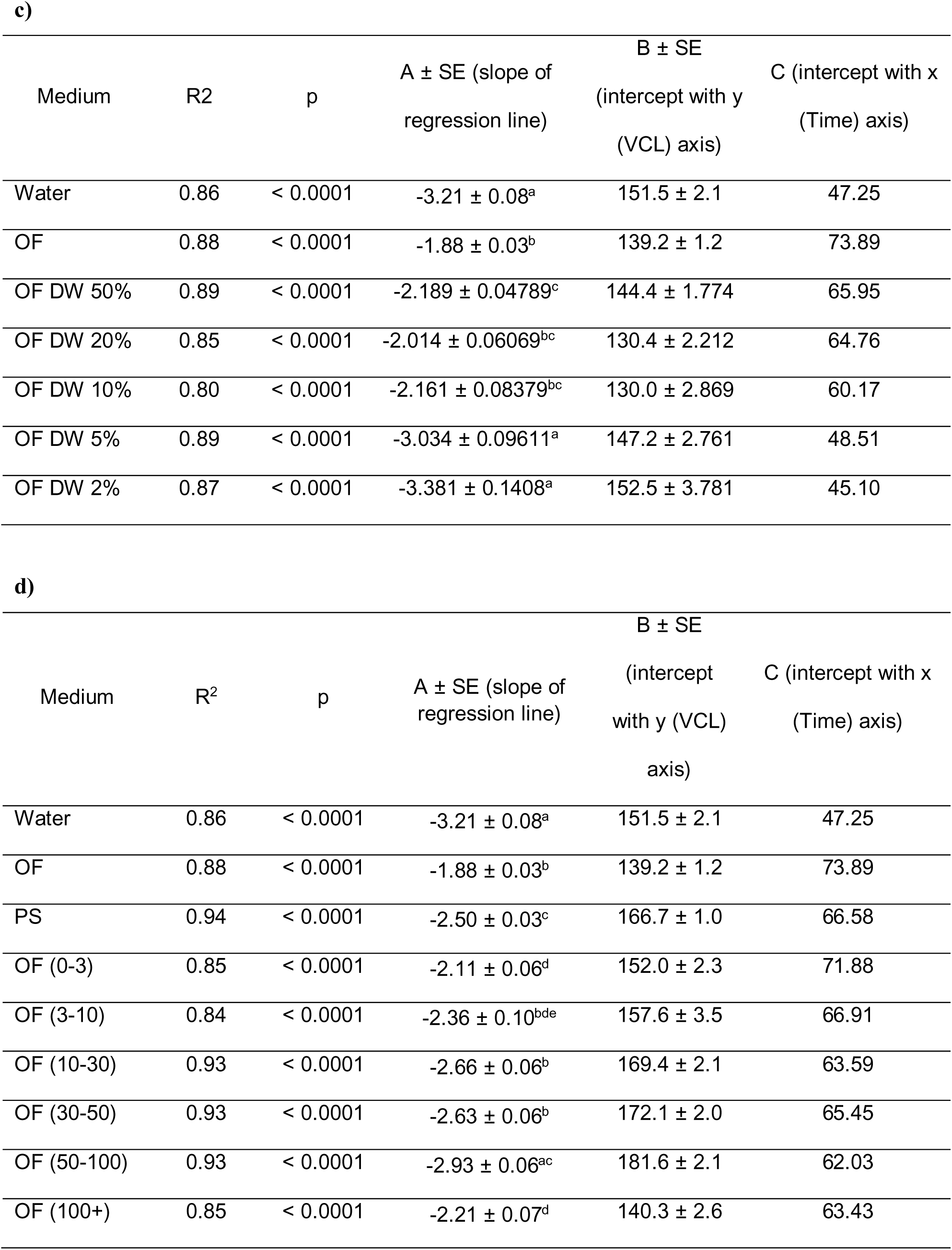
**Parameters of linear regression lines for curvilinear velocity dependencies (Figure 2, Figure 5b) of rainbow trout spermatozoa activated in different media:** distilled water (DW), ovarian fluid (OF); ovarian fluid mixed with distilled water (OF concentration 50; 20; 10; 5 and 2%); NaCl solutions (PS – physiological solution with osmolarity of 290 mOsm/l, diluted with water in the same ratios as ovarian fluid: non-diluted and 50, 20 and 10% of PS in water); and molecular weight fractions of ovarian fluid (with MW 0-3; 3-10; 10-30; 30-50; 50-100 and 100+ kDa). Data A, B and C are mean ± SE, the different superscripts denote significant differences (P < 0.001).

| Medium | R<sup>2</sup> | p | A ± SE (slope of regression line) | B ± SE<br>(intercept with y<br>(VCL) axis) | C (intercept with x<br>(Time) axis) |
| --- | --- | --- | --- | --- | --- |
| Water | 0.86 | < 0.0001 | -3.21 ± 0.08a | 151.5 ± 2.1 | 47.25 |
| OF | 0.88 | < 0.0001 | -1.88 ± 0.03b | 139.2 ± 1.2 | 73.89 |
| PS | 0.94 | < 0.0001 | -2.50 ± 0.03c | 166.7 ± 1.0 | 66.58 |

b)
| Medium | R<sup>2</sup> | p | A ± SE (slope of regression line) | B ± SE<br>(intercept with y<br>(VCL) axis) | C (intercept with x<br>(Time) axis) |
| --- | --- | --- | --- | --- | --- |
| Water | 0.86 | < 0.0001 | -3.21 ± 0.08<sup>a</sup> | 151.5 ± 2.1 | 47.25 |
| PS | 0.94 | < 0.0001 | -2.50 ± 0.03<sup>b</sup> | 166.7 ± 1.0 | 66.58 |
| PS DW 50% | 0.95 | < 0.0001 | -2.723 ± 0.04723<sup>c</sup> | 173.6 ± 1.682 | 63.75 |
| PS DW 20% | 0.90 | < 0.0001 | -3.204 ± 0.08138<sup>b</sup> | 167.9 ± 2.522 | 52.41 |
| PS DW 10% | 0.86 | < 0.0001 | -2.985 ± 0.1002<sup>bc</sup> | 151.5 ± 3.008 | 50.77 |

c)
| Medium | R<sup>2</sup> | p | A ± SE (slope of regression line) | B ± SE<br>(intercept with y<br>(VCL) axis) | C (intercept with x<br>(Time) axis) |
| --- | --- | --- | --- | --- | --- |
| Water | 0.86 | < 0.0001 | -3.21 ± 0.08<sup>a</sup> | 151.5 ± 2.1 | 47.25 |
| OF | 0.88 | < 0.0001 | -1.88 ± 0.03<sup>b</sup> | 139.2 ± 1.2 | 73.89 |
| OF DW 50% | 0.89 | < 0.0001 | -2.189 ± 0.04789<sup>c</sup> | 144.4 ± 1.774 | 65.95 |
| OF DW 20% | 0.85 | < 0.0001 | -2.014 ± 0.06069<sup>bc</sup> | 130.4 ± 2.212 | 64.76 |
| OF DW 10% | 0.80 | < 0.0001 | -2.161 ± 0.08379<sup>bc</sup> | 130.0 ± 2.869 | 60.17 |
| OF DW 5% | 0.89 | < 0.0001 | -3.034 ± 0.09611<sup>a</sup> | 147.2 ± 2.761 | 48.51 |
| OF DW 2% | 0.87 | < 0.0001 | -3.381 ± 0.1408<sup>a</sup> | 152.5 ± 3.781 | 45.10 |

d)
| Medium | R<sup>2</sup> | p | A ± SE (slope of regression line) | B ± SE<br>(intercept with y (VCL)<br>axis) | C (intercept with x<br>(Time) axis) |
| --- | --- | --- | --- | --- | --- |
| Water | 0.86 | < 0.0001 | -3.21 ± 0.08<sup>a</sup> | 151.5 ± 2.1 | 47.25 |
| OF | 0.88 | < 0.0001 | -1.88 ± 0.03<sup>b</sup> | 139.2 ± 1.2 | 73.89 |
| PS | 0.94 | < 0.0001 | -2.50 ± 0.03<sup>c</sup> | 166.7 ± 1.0 | 66.58 |
| OF (0-3) | 0.85 | < 0.0001 | -2.11 ± 0.06<sup>d</sup> | 152.0 ± 2.3 | 71.88 |
| OF (3-10) | 0.84 | < 0.0001 | -2.36 ± 0.10<sup>bde</sup> | 157.6 ± 3.5 | 66.91 |
| OF (10-30) | 0.93 | < 0.0001 | -2.66 ± 0.06<sup>b</sup> | 169.4 ± 2.1 | 63.59 |
| OF (30-50) | 0.93 | < 0.0001 | -2.63 ± 0.06<sup>b</sup> | 172.1 ± 2.0 | 65.45 |
| OF (50-100) | 0.93 | < 0.0001 | -2.93 ± 0.06<sup>ac</sup> | 181.6 ± 2.1 | 62.03 |
| OF (100+) | 0.85 | < 0.0001 | -2.21 ± 0.07<sup>d</sup> | 140.3 ± 2.6 | 63.43 |

**Figure S1.**
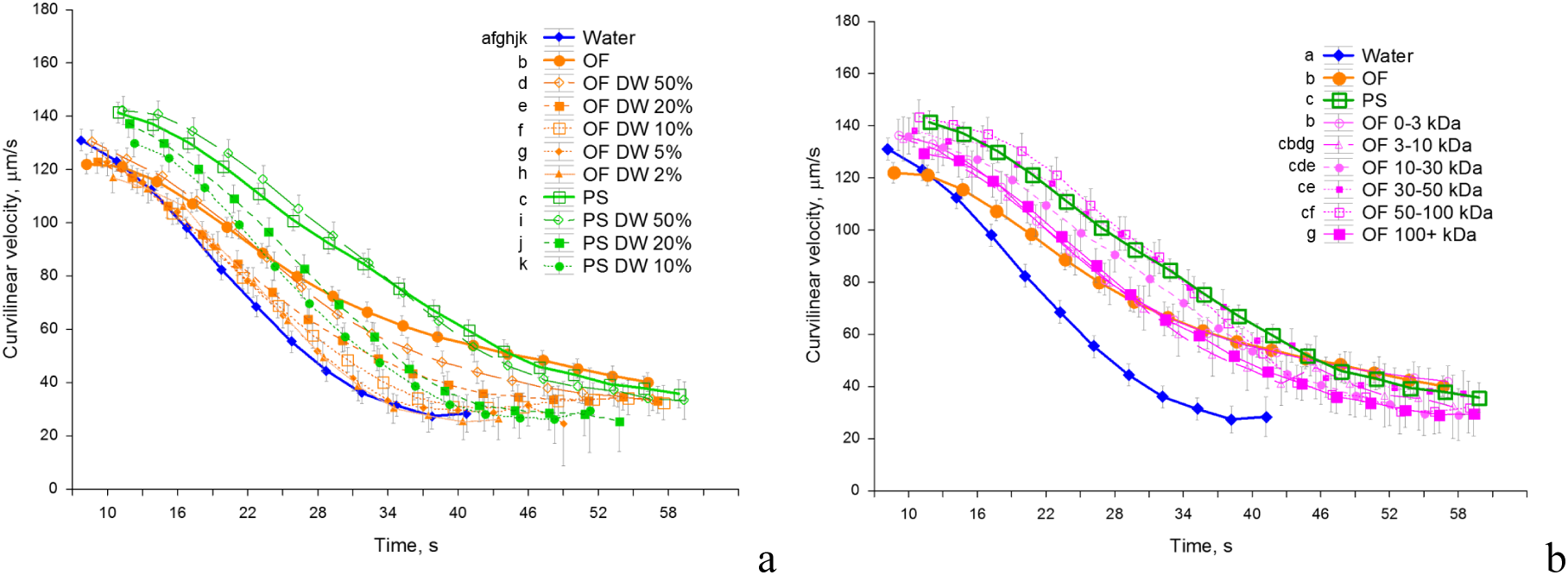
Descriptive statistics data on curvilinear velocity of rainbow trout spermatozoa activated in the presence of ovarian fluid and other conditions, depending on time post-activation: a – velocity in water, ovarian fluid, NaCl solution isotonic to ovarian fluid (physiological solution, PS, 290 mOsm/l); and their dilutions with water (1-1; 1-4; 1-9; 1-19 and 1-49); b – motility in MWCO fractions of ovarian fluid. Data are mean values (average from 5-18 males), vertical bars denote 0.95 confidence interval; different letters in the legends denote significant differences assesses by factorial repeated measures ANOVA with correction for multiple comparisons, p < 0.001. Data are average for 26 males in case of water, ovarian fluid and NaCl isotonic saline; 16 males for OF and 12 for PS dilutions; 11 males in case of MWCO fractions of ovarian fluid.

### Supplementary Video

Response of rainbow trout spermatozoa activated in various media to the introduction of various test fluids using microcapillary: ovarian fluid (OF); distilled water (DW); isotonic NaCl saline (PS, 290 mOsm/l); molecular weight cut-off (MWCO) fractions; thermo-treated (boiled) ovarian fluid; blood serum; distilled water with added 1 mM CaCl_2_. The videos represent periods around 10 s post activation right after activation of spermatozoa and capillary introduction.

https://youtu.be/yqnNJE8gRLc

